# Spatial relationships in the urothelial and head and neck tumor microenvironment predict response to combination immune checkpoint inhibitors

**DOI:** 10.1101/2023.05.25.542236

**Authors:** Alberto Gil-Jimenez, Nick van Dijk, Joris L. Vos, Yoni Lubeck, Maurits L. van Montfoort, Dennis Peters, Erik Hooijberg, Annegien Broeks, Charlotte L. Zuur, Bas van Rhijn, Daniel J. Vis, Michiel S. van der Heijden, Lodewyk F. A. Wessels

## Abstract

Immune checkpoint inhibitors (ICI) currently achieve remarkable clinical results in urothelial cancer (UC). However, the relationship between the tumor microenvironment (TME), usually characterized by immune cell density, and response to ICI is unclear.

We quantified the TME immune cell densities and spatial relationships (SRs) using the multiplex immunofluorescence data of 24 UC pre-treatment tumor resections. We described SRs by approximating the 1-NN distance distribution with a Weibull distribution and evaluated the association between TME metrics (spatial and density parameters) and ipilimumab+nivolumab response.

Immune cell density did not discriminate between response groups. However, the Weibull SR metrics of CD8^+^ T-cells or macrophages to their closest cancer cell were positively associated with response. CD8^+^ T-cells close to B-cells were characteristic of non-response. The G-function, a threshold dependent alternative SR metric, yielded variable effect sizes and statistical power in association studies with response. We validated our SR response associations in a cohort of head and neck tumors with a comparable treatment design. Our data confirm that SRs, in contrast to density metrics, are strong biomarkers of response to ICIs, a finding with significant translational relevance.

## Introduction

Immune checkpoint inhibitors (ICI) block inhibitory signals between immune and neoplastic cells that can result in cancer cell killing^1^. Inhibitors targeting PD-1 and PD-L1 have shown durable responses in a subset of urothelial cancer (UC) patients^2, 3^. Still, most tumors do not respond to treatment^1, 4^, and ICI causes grade≥3 immune-related adverse events in 10% of UC patients^5^. Therefore, it is crucial to identify biomarkers that aid the stratification of responding patients so that alternative lines of treatment can be considered for non-responding patients and prevent unnecessary toxicities.

The surrounding of a tumor, known as the tumor microenvironment (TME), contains immune cells, normal epithelial cells, and fibroblasts that continuously interact^6^. Components of the TME indicative of pre-existing immunity have shown associations with response to ICI, such as CD8^+^ T-cell infiltration and transcription factors related to T-cell activity^7–11^. However, biomarkers do not behave consistently in UC trials. For instance, the baseline presence of CD8^+^ T-cells correlates with ICI monotherapy response in the neoadjuvant (anti-PD-L1)^7^ or metastatic setting^12^. Still, in pre-operative ICI combination therapy (anti-PD(L)-1 + anti-CTLA-4), the response is independent of CD8^+^ T-cell density^9, 10^. The lack of robust response biomarkers highlights the need to dissect tumor-immune interactions in more detail^13^.

A technology enabling a TME characterization at single-cell resolution is multiplex immunofluorescence (mIF), which spatially profiles a tissue slide using multiple antibodies simultaneously^14, 15^. MIF-derived metrics were found to predict anti-PD-1 and anti-PD-L1 response across different tumor types^16^, highlighting its ability to quantify crucial immune components that determine ICI response. Typically, mIF data are summarized as immune cell densities, informing about immune cell counts, and typically topologically assessed in different compartments, i.e., tumor and stroma^17^.

By definition, immune cell density and abundance metrics ignore the immune interactions relevant to an anti-tumor response^18^. Ignoring these interactions is suboptimal, as many immune interactions require proximity. For instance, a T-cell receptor and antigen interaction require physical binding, which requires the cells involved to be in close proximity to each other. In contrast, an immunosuppressive TME will have few immune cells close to cancer cells due to their inability to infiltrate a tumor^19^. Such distance or adjacency patterns between cells at the TME can be measured through their spatial relationship (SR), allowing for a mathematical description amenable to downstream analysis. Notably, associations between SRs at the TME and prognosis^18, 20, 21^ and response to monotherapy ICIs have been reported across different cancer types^22–24^. The SRs allow for quantitative exploration of the TME, providing a basis for improving our understanding of tumor immunology and scrutinizing new associations with ICI response. SRs in the UC’s TME are poorly understood and, to our knowledge, largely unexplored in neoadjuvant combination ICI treatments, and this study aims to address that.

Several analytical frameworks aim to measure the SRs of the TME, such as cell-cell interactions and tissue modules, using spatially-resolved protein-derived data across multiple data types^19^. Methodologies are predominantly topologically based (graph-, networks-, and cell-counting-based methods) or distance-based. Distance-based methods, such as the first-nearest-neighbor (1-NN) distribution, allow modeling spatially-resolved data. Because distances following a 1-NN distribution are asymmetrical, approaches that estimate the 1-NN distribution mean^24^ provide inadequate data summaries. A common approach to model 1-NN distributions is through the cumulative distribution function (CDF), known as the G-function. Nevertheless, the downstream analyses require an additional summary by estimating the area under the curve (AUC) at a predefined threshold^25, 26^. Currently, there is a lack of spatial methodologies that describe the distance distribution without using a threshold and that model variation between individuals.

In this study, we spatially profile cancer cells, T-cells, macrophages, and B-cells using mIF and present a methodology to quantify the TME SRs using 24 pre-treatment UC samples from the NABUCCO trial^10^. In NABUCCO, pre-operative combination ICI with ipilimumab and nivolumab is administered in UC. We fit a Weibull distribution to the 1-NN distance distribution between pairwise cell relationships to extract a two-parameter describing the distribution. These descriptions outperformed immune cell densities when quantifying the differences in immune cell SRs between response groups to ICI. To demonstrate generalizability of our findings, we confirmed the baseline associations between SRs and response in an independent cohort of 25 mostly HPV-negative head and neck squamous carcinoma (HNSCC) patients receiving neoadjuvant ipilimumab and nivolumab treatment^27^.

## Results

### Multiplex immunofluorescence and modeling of immune cell densities and spatial relationships of urothelial and head and neck cancer samples

We collected multiplex immunofluorescence (mIF) data from baseline formalin-fixed, paraffin-embedded (FFPE) stage-III urothelial cancer (UC) samples (n=24), of patients recruited in the NABUCCO trial (**Figure 1A**). Patients received pre-operative combination ICI, consisting of two or three cycles of ipilimumab (anti-CTLA-4) and nivolumab (anti-PD-1)^10^. We determined the position and identity of cells using mIF and identified B-cells, T-cells (CD8^+^ T-cells, FoxP3^+^ T-cells, and T-helper cells), macrophages, and cancer cells (**Figure 1B**). Negative cells scored negative for all the antibodies (CD8^-^, CD3^-^, FoxP3^-^, CD20^-^, CD68^-^, PanCK^-^); this group contains all stromal cells and immune cells not covered by our antibody panel. Next, by comparing the local density of cancer and negative cells, we virtually segmented the tissue into the tumor and stroma compartments (**Figure 1C, Supplementary Figure 1**) and quantified immune cell density in both compartments (**Figure 1D**, **Supplementary Table 1**).

**Figure 1.**
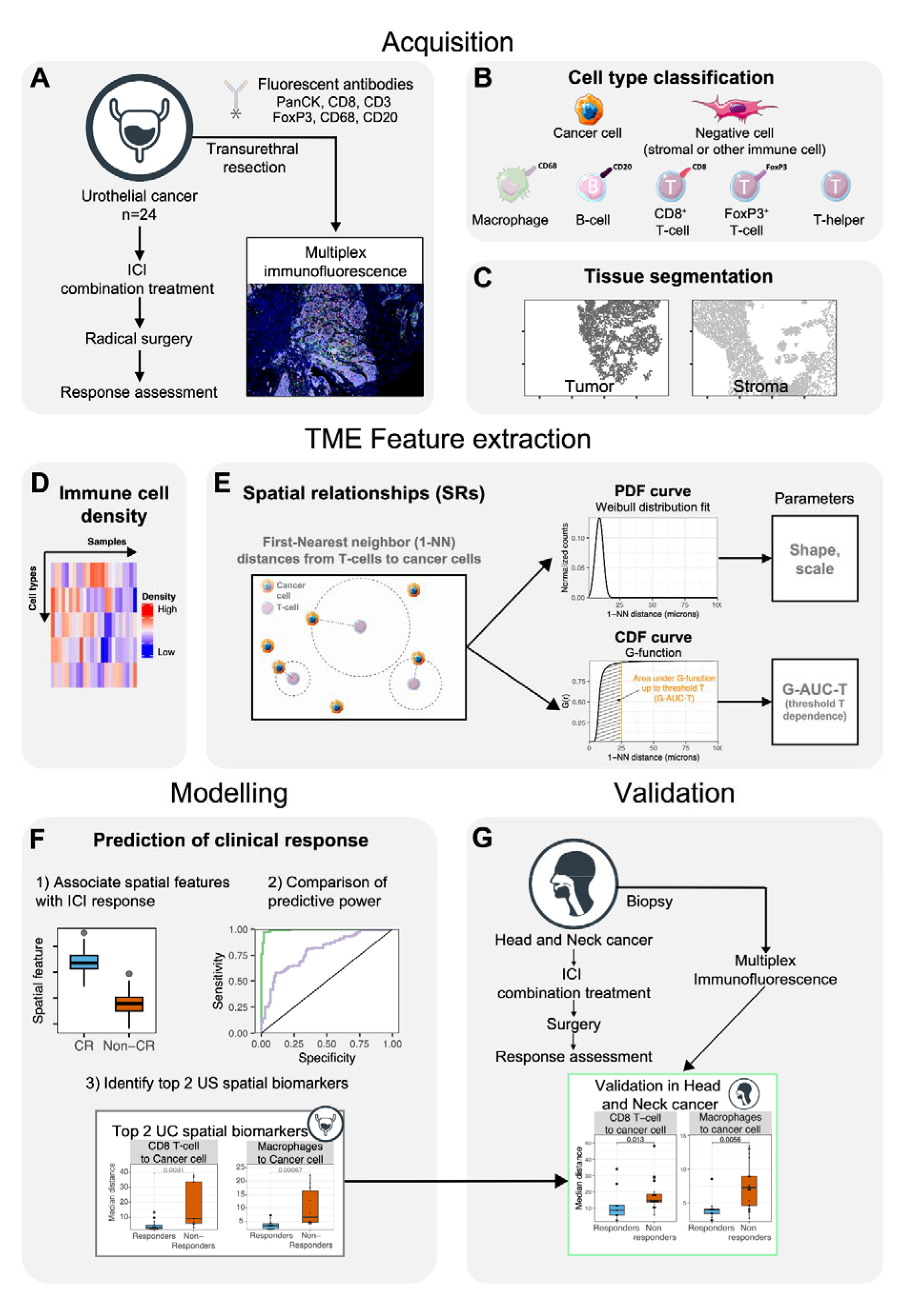
Profiling of immune cell density and spatial relationships of the urothelial cancer tumor micro-environment by multiplex immunofluorescence. (A) Biopsy samples from 24 patients from the NABUCCO trial were profiled using mIF. (B) Cell type classification by comparing antibody marker positivity. (C) Tissue segmentation into tumor and stroma regions by comparing the local densities of cancer cell marker positive and negative cells. (D) Immune cell density in the tumor and stroma compartments was calculated in each tissue compartment (tumor and stroma). (E) SRs were summarized using the 1-NN statistic studied from a reference cell type to a target cell type. The resulting 1-NN distances vector was studied using 2 approaches: modeling a Weibull distribution to the Probabilistic Density Function (PDF) (top), and using the cumulative distribution function (CDF) using the G-function (F) Association of SR parameters with ICI response and comparison of the discriminative power between SR and density TME parameters. (G) Validation of associations between SR parameters and response identified in UC in an independent cohort of HNSCC tumors. Abbreviations*: TME*: tumor micro-environment; *SR*: spatial relationship; *mIF*: multiplex immunofluorescence; *ICI*: immune checkpoint inhibitors; *1-NN*: first nearest neighbor; *PDF*: probabilistic density function; *CDF*: cumulative density function; *G-AUC-T*: G-function evaluated at a threshold T; *T*: threshold; *UC:* urothelial cancer; *HNSCC*: head and neck squamous cell carcinoma.

We quantified the pairwise SRs between all cell types in the TME using the first nearest-neighbor (1-NN) distance statistic (**Supplementary Table 2**). In brief, the statistic is measured between a reference cell type (*cell from*) and a target cell type (*cell to*). The distances between each reference cell type and their closest target cell type yielded a 1-NN distance vector (**Figure 1E**). Then, we fitted a Weibull distribution function using a non-linear mixed effect model to summarize the 1-NN distance distribution. The model has two parameters: shape and scale, which uniquely describe the 1-NN distance distribution (**Figure 1E**, top) in a threshold-independent manner. We estimated the Weibull parameters (scale and shape) for all 49 pairwise relationships between cell types for all samples using the data from the whole tissue slide. Additionally, we evaluated the G-function derived from the cumulative distribution function (CDF) of the 1-NN distance distribution and broadly documented in the spatial statistics literature^28^. The G-functions were summarized by computing the area under the curve (AUC) at different thresholds T, which we refer to as G-AUC-T (which included T = 25 [**Figure 1E**, bottom], T = 50- and T = 100-micron) across all pairwise cell type SRs and samples.

We then compared the spatial (**Figure 1E**) and density (**Figure 1D**) parameters with response to ICIs and compared their predictive power (**Figure 1F**). Lastly, we validated the associations between TME parameters and response in an independent cohort of mostly HPV-negative head and neck squamous cell carcinoma (HNSCC, n=25) using baseline primary tumor samples of patients recruited in the IMCISION trial that received pre-operative ipilimumab and nivolumab combination treatment (**Figure 1G**).

### Exploration of spatial relationships across the urothelial cancer tumor micro-environment

We quantified the SRs for all pairwise relationships of immune, cancer, and negative cells by estimating the Weibull parameters (shape, scale) characterizing the 1-NN distance distributions (**Figure 1E**). Next, we explored the shape-scale parameter space across patients (**Figure 2A**).

**Figure 2.**
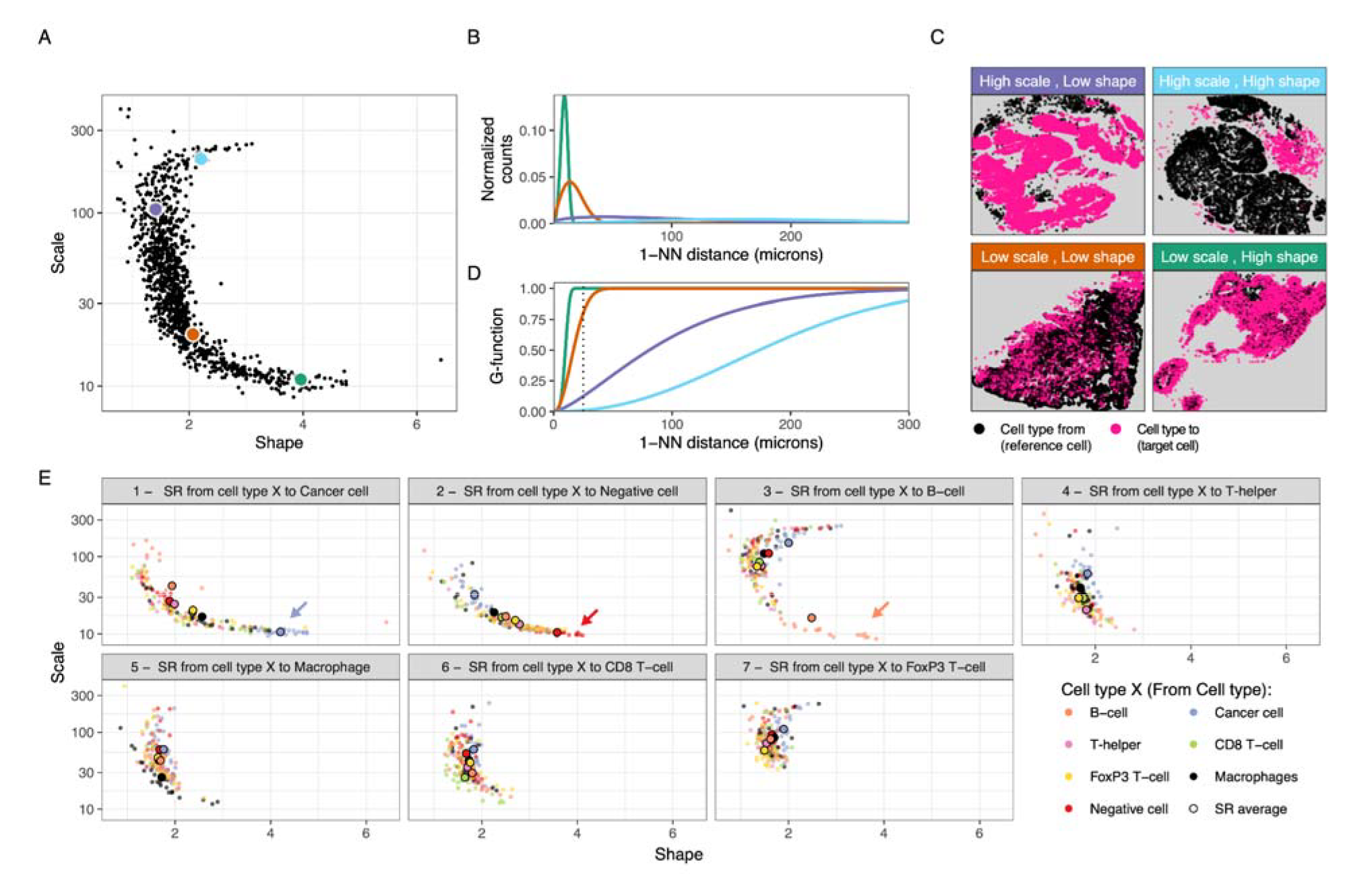
Exploration of pairwise cell type SRs in the TME using the 1-NN distance statistic. (A) Scale vs shape SR parameters fitted on the 1-NN distribution for the 24 samples and the 7×7 cell type combinations. Representative examples are highlighted in green, orange, blue and pink, with their associated 1-NN distance distributions (B), point patterns (C) and G-functions (D). (E) Scatter plot of the scale-shape parameter space by target cell type (cell type to in the SR). For instance, the first facet represents SRs studied from any reference cell type to cancer cells. There, the coloring denotes the reference cell type (cell type from); i.e. the orange dots represent spatial relations studied from B-cells to cancer cells. Cohort averages for their associated SR parameters are highlighted as big dots for each cell type-cell type combination. Abbreviations: *SR*: spatial relationship; *TME*: Tumor microenvironment; *1-NN*: First nearest neighbor.

To illustrate what the Weibull parameters (shape, scale) represent, we use four combinations of the SR metrics that characterize distinct instances of cellular spatial distributions (**Figure 2A**, colored dots). For the green dot in **Figure 2A**, we observe that the 1-NN distance distribution is characterized by low distances (**Figure 2B**, green curve), which originate from the black cells being close to a pink cell (**Figure 2C**, “Low scale, High shape”). The low scale/high shape value signals densely packed pink cells, meaning there is always a pink cell close to a black cell. Another type of SR characterized by relatively short 1-NN distances but with a higher variance in 1-NN distances, is illustrated in **Figures 2A-B** (orange dot and curve). It represents an SR where one or both cell types are arranged in overlapping, densely packed clusters, such as the pink and black cells in **Figure 2C** (*“Low scale, Low shape”*). Cases with an even larger spread in distances (**Figures 2A-B**, purple dot and curve) from the black to the nearest pink cells, are described by high scale and low shape parameter values (**Figure 2C**, *“High scale, Low shape”*). Lastly, the cyan dot (**Figure 2A**) represents a 1-NN distribution shifted towards high 1-NN distances (**Figure 2B**, cyan curve), characteristic of a repulsion pattern, i.e., where both cell types are clustered in relatively large clusters (**Figure 2C**, *“High scale, high shape”*). Furthermore, the associated G-function results showed corresponding differences between the four scenarios, which is expected, as the G-function is the cumulative distribution of the 1-NN distances (**Figure 2D**, **equation 4**). However, in contrast to the Weibull approach, a threshold value (T) is required to generate the summary metric G-AUC-T.

We then dissected the SRs by reference and target cell type to explore patterns of SRs across the TME (**Figure 2E**). First, we investigated self-self relationships, which are relationships between cells of the same type. The self-self relationship for tumor cells falls in the “*Low scale, High shape*” scenario, with short distances between cells as tumor cells are typically densely packed in the tumor regions (**Figure 2E**-1, blue dots and arrow). Similar behavior was observed for negative cells, indicating a tight packing of negative cells (**Figure 2E**-2, red dots and arrow). Then, we explored the self-self relationships of immune cells. We observed B-cells clustering for all patients (**Figure 2E**-3, orange dots and arrow). We did not observe the same clustering behavior for self-self SRs of other immune cell types (e.g., green dots in **Figure 2E**-6, which shows a behavior more akin to the *“Low scale, Low shape”* scenario). Subsequently, we assessed the SRs between different cell types. We observed a high variation in the Weibull parameters across samples and pairwise cell type combinations (**Figure 2E**) and a dependence on the SR “perspective”, i.e., whether a given cell type is the cell type from or the cell type to.

**Figure 3.**
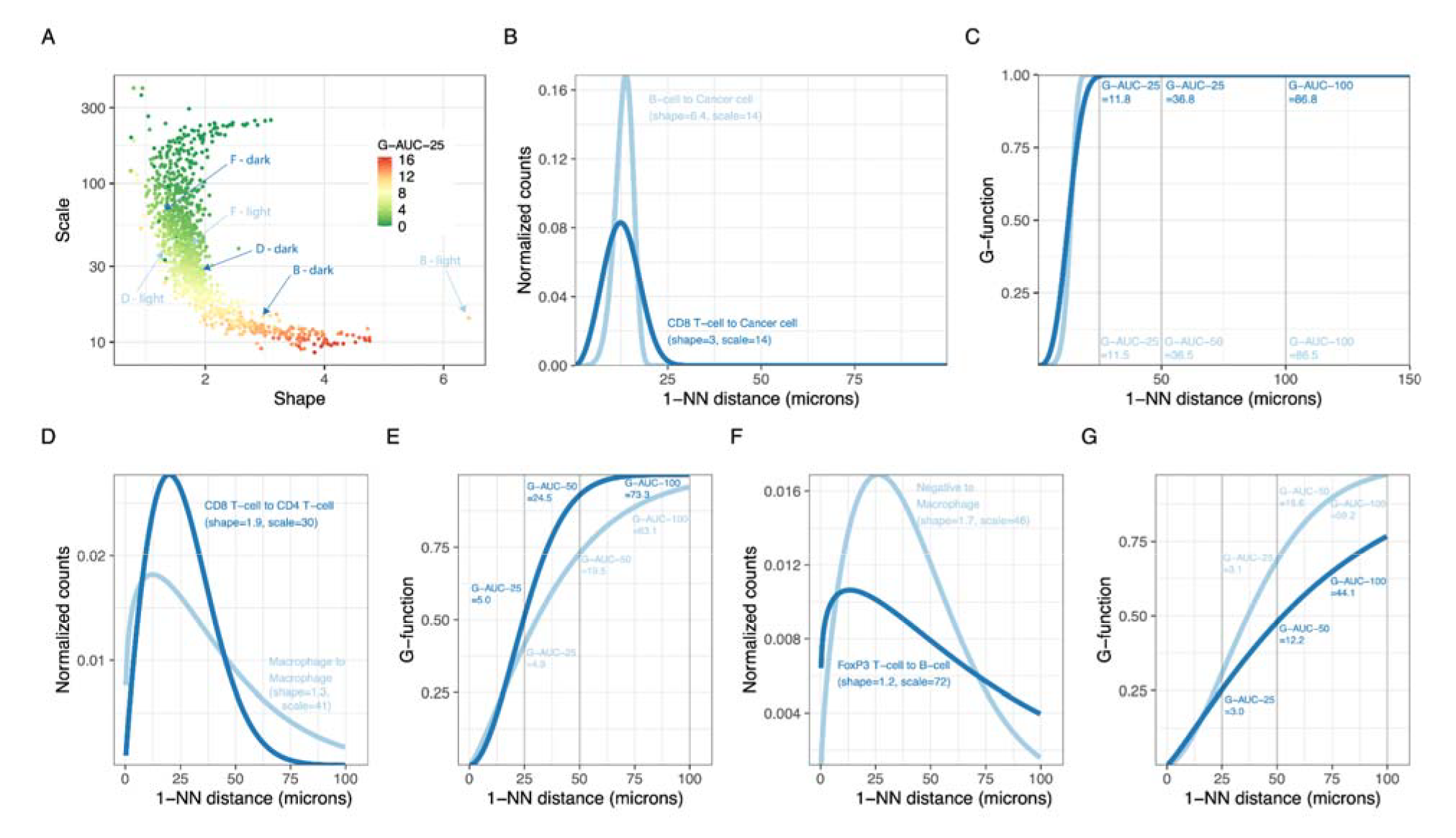
Comparison of spatial relationship parameters derived from the 1-NN distance statistic. (A) Scale/shape parameter space associated with the G-AUC-25 (coloring, G-function AUC evaluated at 25 microns). (B, D, F) 1-NN PDF distance distribution of pairs of examples with a substantial difference in the shape parameters. The plot reports each curve’s shape, scale, and SR. Each SR is annotated in panel (A). (C, E, G) Associated G-functions (CDF) of the examples shown in panel A of pairs of examples without a substantial difference in G-AUC-25. G-AUCs evaluated at different thresholds T are reported in the figure. Abbreviations: *AUC*: Area under the curve; *G-AUC-T*: G-function AUC evaluated at a threshold T; *T*: threshold; *1-NN*: first nearest neighbor; *SR*: spatial relationship.

In short, we created a framework to quantify, interpret, and study SRs in the TME using the Weibull parameters extracted from the 1-NN distance distributions. The framework allows exploring distinct cellular organization patterns and quantifying specific SRs (e.g., T-cell to B-cell vs. B-cell to T-cell) amenable for downstream analyses aimed at furthering our understanding of the TME and its relationship to response to ICIs.

### Comparison between spatial relationship metrics derived from the first nearest neighbor distance statistic

A common approach to extracting parameters from the 1-NN distance distribution statistic is through the G-function, which represents the cumulative density function (CDF) of the 1-NN distribution and requires a particular threshold to summarize the data for downstream analyses. We compared our Weibull parameters with the G-function summary. We evaluated the G-function using its area under the curve (AUC) up to a 25-micron distance, which we defined as the “G-AUC-25”. We chose this threshold after inspecting the distance region in which the G-function showed the highest variability. Because we observed variability in G-functions across pairwise cell-type relationships, other thresholds were evaluated and denoted as “G-AUC-T”, in which T denotes the threshold in microns. We observed a non-linear relationship between the shape, scale and the G-AUC-25 (**Figure 3A**), G-AUC-12 and G-AUC-50 (**Supplementary Figures 2A-B**).

Upon summarizing the G-function, and for a given SR’s G-AUC-25 value, we observed that the associated shape and scale parameters can show substantial variation (**Figure 3A;** e.g., dots coloured in red mapping to a wide range of shape parameter values). **Figure 3A** highlights a pair of SRs with large differences between their shape or scale parameters, which can be visually confirmed by their 1-NN curves (**Figures 3B**, **D**, **F**). In contrast to the Weibull parameters, the corresponding G-AUC-25 values do not differ substantially between the pair members (**Figures 3C**, **3E**, and **3G**, showing the G-function curves corresponding to the pairs in **Figures 3B**, **3D,** and **3F,** respectively), which can hinder interpretations on the associated SR. Specifically, comparing **Figures 3B** and **3C,** we observe that while the Weibull parameters are quite different (scale=14 and shape=6.4 for B-light, scale=14 and shape=3.0 for B-dark) the G-AUC-T values are quite similar (G-AUC-25=11.5 for B-light and G-AUC-25=11.8 for B-dark). While in the other examples the differences in the G-AUC-25 values were small, we found that higher values of the summary threshold could better capture the difference between the SR pairs in **Figure 3E** (Weibull density in **3D**) and **Figure 3G** (Weibull density in **3F**).

To further illustrate differences between SRs captured by the shape or scale parameters but not by the G-AUC-25 parameter, we compared G-AUC-Ts for different pairwise cell-cell relationships (**Supplementary Figure 2C**). Here, for different samples but the same SRs (e.g., *Macrophages to B-cells*), the magnitude of the increase in the G-AUC-T value when altering the evaluation threshold T, depending on the studied pairwise cell-cell relationship, the G-function’s shape (rapidly or slowly reaching the maximum value), and the sample. In some SRs, the G-AUC’s increase was linear (e.g., *Cancer cell to cancer cell*) because the G-function saturated at low thresholds (**Supplementary Figure 2D**). Still, in others (e.g., *Macrophages to B-cell*), the increase was not always linear because the G-function reaches saturation at higher thresholds (**Supplementary Figure 2D**). Therefore, when using a G-function statistic, such as the G-AUC-T, the SR quantification critically depends on the threshold used.

In short, our data shows that the G-function threshold introduces variance in the downstream metric G-AUC-T. Furthermore, optimizing the threshold to maximize the effect size of SR-biomarkers for treatment response creates a risk of overfitting. Therefore, different results using the same SR data can be obtained when varying the threshold, which can hinder downstream interpretation.

### Spatial relationships associated with immune checkpoint blockade response

Multiplex immunofluorescence data are usually summarized as cell type fractions or immune cell density. We quantified the density of T-cells, B-cells, and macrophages in both the tumor and stromal compartments. However, we found no significant differences between response groups (**Figure 4A**), indicating no differences between response groups in immune cell abundances in either the tumor and stromal compartments. In addition, immune cells spatially distribute following configurations of immune phenotypes^29^, being *Excluded* (high immune cell abundance in the stroma), *Inflamed* (high immune cell abundance in the tumor), and *Desert* (low immune cell abundance in the tumor and stroma). We quantified exclusion ratios (ratio between stromal and intratumoral immune cell density) and used them as a proxy of immune phenotypes for each immune cell. However, again we found no significant associations with treatment response (**Figure 4B**), suggesting similar immune cell configurations in the response groups.

**Figure 4.**
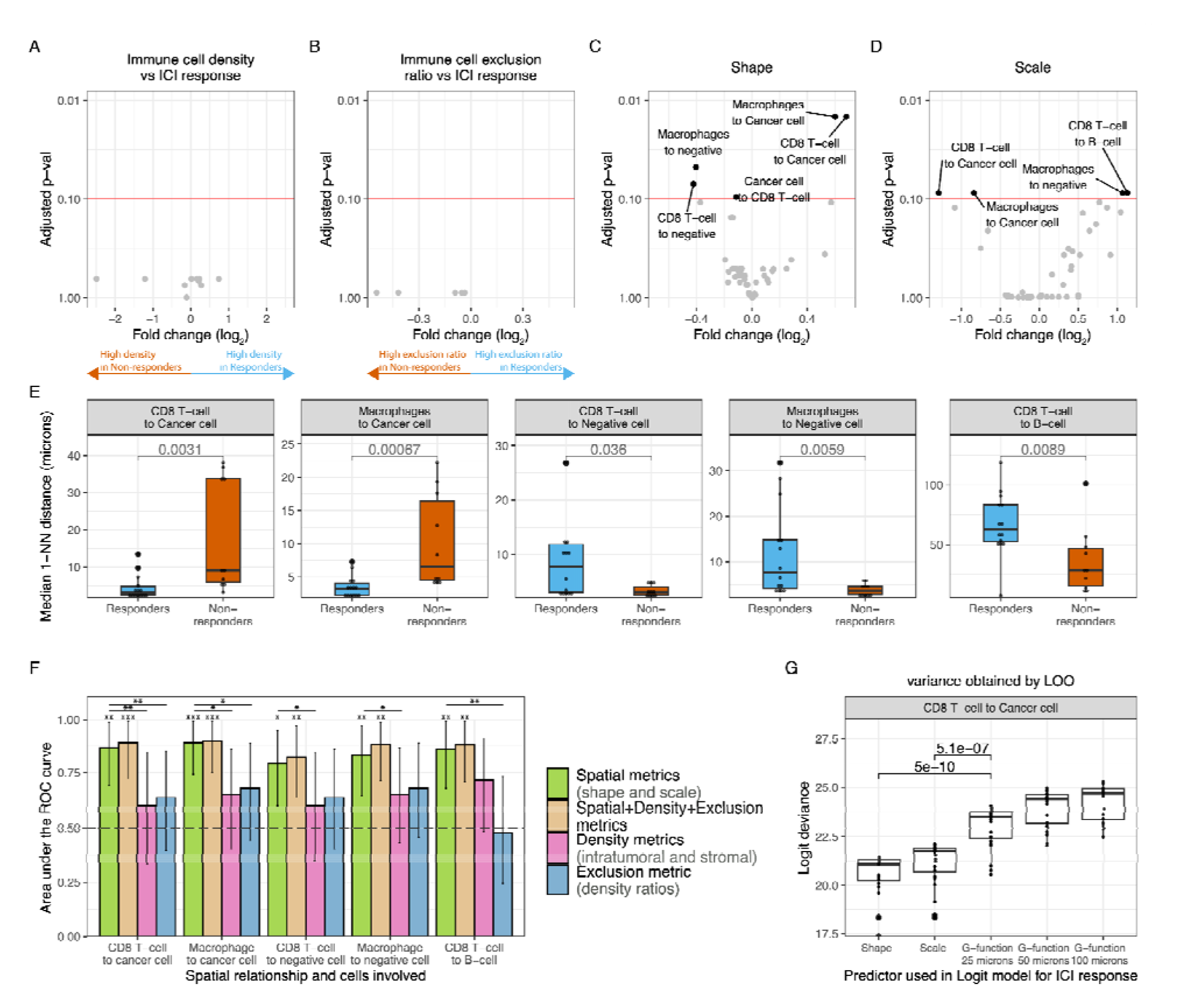
Association of spatial relationships with response to pre-operative ipilimumab+nivolumab in urothelial cancer. (A) Volcano plot showing the fold change on the intratumoral and stromal immune cell densities between response groups (x-axis) and statistical significance by t-test adjusted by multiple hypothesis testing (y-axis). (B) Volcano plot showing the fold change on the exclusion ratio (ratio between stromal and intratumoral immune cell density) between response groups (x-axis) and statistical significance by t-test adjusted by multiple hypothesis testing (y-axis). (C) Volcano plot showing the fold change on the shape parameter between response groups (x-axis) and statistical significance by t-test adjusted by multiple hypothesis testing (y-axis). (D) Volcano plot showing the fold change on the scale parameter between response groups (x-axis) and statistical significance by t-test adjusted by multiple hypothesis testing (y-axis). (E) Median first nearest-neighbor distance distribution per response group as calculated by the associated shape and scale for the SRs that are significantly associated with response to ICI treatment and statistical significance by a Mann-Whitney test. (F) ROC curve AUCs of the discriminative power of distinct TME parameters: all spatial parameters (green), all spatial parameters, density and exclusion ratio metrics (gold), all density metrics (pink), and exclusion ratio between stroma and tumor density (blue). The lines denote whether a statistical significance on the associated AUC was achieved. Significance symbols above bar plots denote whether the AUC is significantly different from AUC=0.5. Significance symbols between bar plots denote whether the ROC-AUC-spatial (green) is significantly greater than the ROC-AUC-spatial-density-exclusion (gold) or ROC-AUC-density (pink) or ROC-AUC-exclusion (blue) by bootstrapping 500 times. Confidence intervals of ROCs were estimated by bootstrapping samples 500 times. (G) Logistic regression deviance of a univariate logistic regression model predicting ICI response using as a predictor the shape, the scale, or the G-function evaluated at different thresholds (G-AUC-T). Variability on the AIC was evaluated by leave-one-out cross-validation and significance was tested by a student’s t-test. Unless otherwise stated, all statistical tests were two-sided. Significance symbols: *: p<0.05, **: p<0.01, ***: p<0.001. Abbreviations: *ICI*: Immune checkpoint inhibitor; *ROC*: Receiver operating characteristic; *AUC*: area under the curve; *TME*: tumor microenvironment; *G-AUC-T*: G-function evaluated at a threshold T; *T*: threshold; *logit*: logistic regression.

Motivated by the invariance between immune cell abundances and density ratios in the TME between response groups, we investigated whether spatial relationships derived from the TME were predictive of response to ICI combination treatment. We first investigated whether the SRs of all pairwise cell types, characterized by the Weibull parameters (shape and scale), were associated with clinical response (**Figure 1F**). After correction for multiple hypothesis testing, we identified nine SRs that were associated with clinical response (FDR<0.10) for the Weibull parameters shape and scale (**Figures 4C, D**). The association between G-AUC-T and response using a rank-based statistic strongly depended on the selected value of the threshold, with the fold change decreasing with increasing values of the threshold (T) (**Supplementary Figure 3**). When selecting a low threshold value (T = 25 microns), we found no significant associations between SRs quantified by G-AUC-25 and response. Upon increasing the threshold value (T = 50 microns), we found three associations between SRs quantified by G-AUC-50 and response, of which two were also identified using the Weibull parameters and one SR (*FoxP3^+^ T-cell to negative cell*) was trending but not significant (FDR_scale_=0.21, FDR_shape_=0.11) using the Weibull parameters.

To guide interpretation, we computed, for each SR significantly associated with clinical response to combination immunotherapy and each patient, the median 1-NN distances (**Figure 4E**). In responding tumors, the distances from either CD8^+^ T-cells or macrophages to the closest cancer cells were smaller than in non-responders (median 1-NN distance *CD8*^+^ *T-cell to cancer cell*, responders=4±3μm, non-responders=18±15μm; *Macrophage to cancer cell*, responders=4±2μm, non-responders=10.2±7μm). Conversely, responding tumors had the largest 1-NN distances for the SR from CD8^+^ T-cells or macrophages to the closest negative cell (median 1-NN distance *CD8*^+^ *T-cell to negative cell*, responders=9±8μm, non-responders=3±1μm; *Macrophage to negative cell*, median 1-NN distance responders=12±10μm, non-responders=4±1μm). Despite the clear differences in the associated median 1-NN distances, the G-function approach did not identify the associations of the SRs with response at a low threshold (T = 25 microns, **Supplementary Figure 3A**) nor the associations of the SRs involving CD8^+^ T-cells and response at a higher threshold (T = 50 microns, **Supplementary Figure 3B**, FDR=0.14 and FDR=0.20 for *CD8*^+^ *T-cell to cancer cell* and *CD8*^+^ *T-cell to negative cell*, respectively). Furthermore, non-responding tumors were characterized by small distances from CD8^+^ T-cells to B-cells (median 1-NN distance *CD8*^+^ *T-cell to B-cell*, responders=66±27μm, non-responders=36±27μm). We identified an association between the SR from cancer cells to CD8^+^ T-cells and response with a small fold change for the shape parameter (|FC_shape_|=0.11, FDR_shape_=0.09), pointing to a difference in distribution that was not detected in terms of median 1-NN distances (**Supplementary Figures 3A-E**, p=0.9), suggesting that this may well be a false positive. Lastly, the single SR biomarker identified by the G-function approach at a 50 microns but not at a 25 microns threshold (*FoxP3 T-cell to negative cell*, FDR_G-AUC-25_=0.12, FDR_G-AUC-50_=0.04, **Supplementary Figure 3** and **Supplementary Figure 4J**) that was not identified by the Weibull approach was trending (**Supplementary Figure 4I**, FDR_shape_=0.11) and showed relative differences in the associated median 1-NN distances (**Supplementary Figure 4F**).

In contrast to the SRs, the immune cell density and exclusion ratios were not associated with response. We confirmed, using simulated data, that density affects SRs between rare (e.g., immune to immune cells) but not between abundant and rare cell types (e.g., cancer to immune cells) (**Supplementary Note 1**). To further confirm independence between density and SR metrics in the predictive setting, we compared the predictive power for clinical response of the SR Weibull parameters (shape and scale) and their associated relevant density and exclusion metrics. The comparisons were made for each SR that was significantly associated with treatment response. For example, the SR from CD8^+^ T-cells to cancer cells was associated with response (FDR_shape_= 0.01, FDR_scale_=0.09). We compared its predictive power with the CD8^+^ T-cell density (intratumoral and stromal) and exclusion ratio of CD8^+^ T-cells. For this comparison, we employed a logistic regression model and the resulting area under the ROC curve (AUROC). The AUROCs for the five SR associations (depicted in **Figure 4E**) and the associated density and exclusion metrics are shown in **Figure 4F**. No density or exclusion metric reached significance as all 95% CI of the associated AUROCs included AUROC=0.5. In contrast, all the SR Weibull parameters reached significance with AUROC values around 0.8 and 95% CI that do not include AUROC=0.5 (**Table 1**, **Supplementary Figure 5**), highlighting the superior predictive power of the SR metrics. Lastly, we tested whether adding density and exclusion ratio as covariates to the SR-based model improved the performance, but this was not the case (**Figure 4F**, **Table 1**).

**Table 1.**
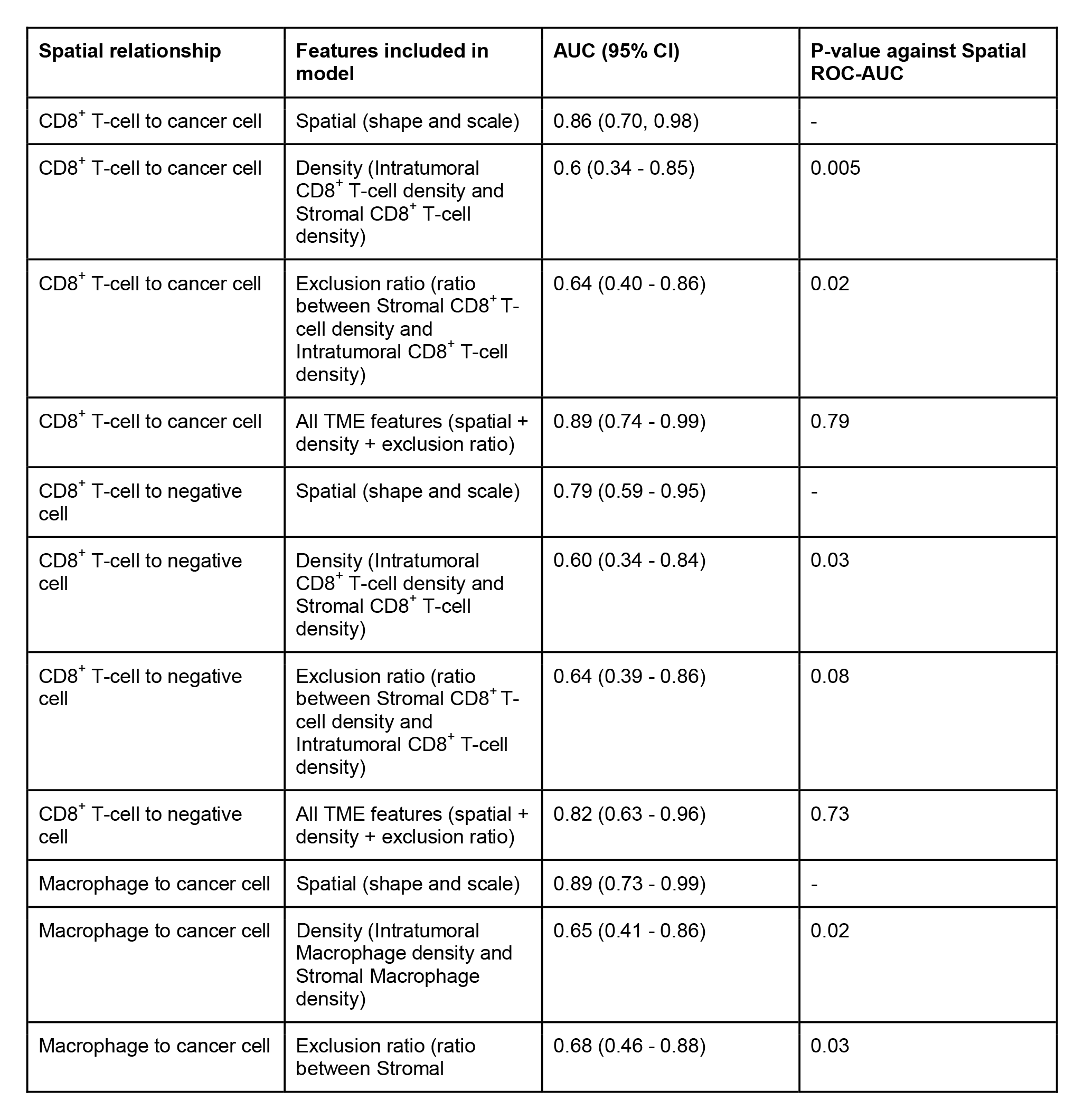
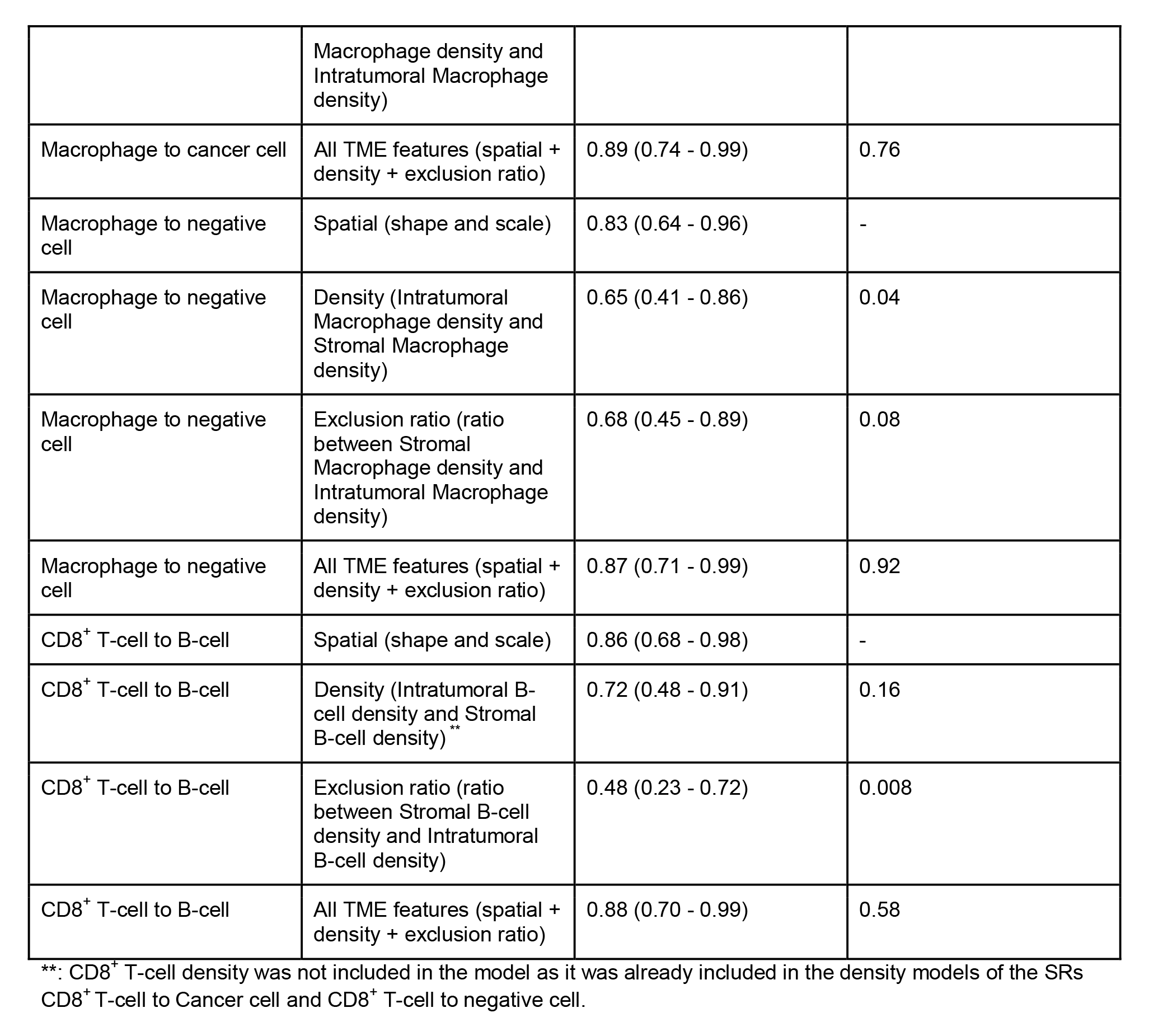
Predictive power of logistic regression models as measured by the ROC-AUC for logistic regression models trained using only spatial metrics (shape, scale), density metrics (intratumoral and stromal density), exclusion ratio metrics (ratio between stromal and intratumoral density), and all metrics (all the metrics above). P-value against spatial ROC-AUC denotes the difference between each model and the model trained spatial metrics from the associated SR by bootstrapping 500 times (testing whether the AUC is greater). Confidence intervals of ROCs were estimated by bootstrapping samples 500 times. Abbreviations: ROC: receiver operating characteristic; AUC: area under the curve; SR: spatial relationship.

Next, we compared different SR metrics derived from the 1-NN distance distribution (shape, scale, G-AUC-T at different T) for their ability to describe clinical response. To do so, we compared the Weibull parameters to the G-AUC-T metrics in the predictive setting. Specifically, we compared the logistic regression deviance, in which lower values indicate better model fits. We did so for the six SRs that were found to be significantly associated with response. The SR from CD8^+^ T-cells to cancer cells model showed that the shape or the scale parameters scored significantly better than the G-AUC-T trained models for T = 25, 50, and 100 (**Figure 4G**). For the remaining SRs significantly associated with clinical response (**Figures 4C-D**), we observed that the models trained using Weibull parameters (shape and scale) outperformed the models trained using G-function parameters, except for the SR from macrophages to negative cells and from CD8 T-cells to B-cells where G-AUC-25 performed similarly as in the Weibull approach (**Supplementary Figure 6**).

In summary, we observed that mIF-derived spatial relationships in the TME hold superior predictive power for clinical response compared to immune cell density or immune phenotypes. Our results convincingly demonstrate that the Weibull parameters (shape and scale) are superior to the G-function metrics (G-AUC-T) in predicting clinical response to combination checkpoint therapy.

### Validation of spatial relationships biomarkers of ICI response in a cohort of head and neck cancer

We tested whether our spatial biomarkers also predicted response in other cancer types. We used a cohort of head and neck squamous cell carcinoma (HNSCC) patients from the IMCISION trial^27^ to validate our findings. A subset of 25 IMCISION patients was treated with pre-operative ipilimumab+nivolumab (similar to NABUCCO), and successfully provided tumor sample profiling with the same mIF antibody panel as the UC cohort (**Figure 1G**).

We first compared the SR parameter space in HNSCC (**Supplementary Table 3**) with that of the UC cohort. We observe the same “C-shape” distribution in the shape-scale space as we observed in UC (**Figure 5A**). Second, we found a high concordance between NABUCCO and IMCISION for the shape and scale population averages across pairwise SRs between all cell types (**Figure 5B**, **Supplementary Figures 6A-D**). These results suggest that the distance between cell types follows a characteristic pattern preserved across these two cancer types. For instance, similar behavior of B-cell to B-cell 1-NN distances compatible with the “*Low scale, High shape*” behavior was observed in HNSCC.

**Figure 5.**
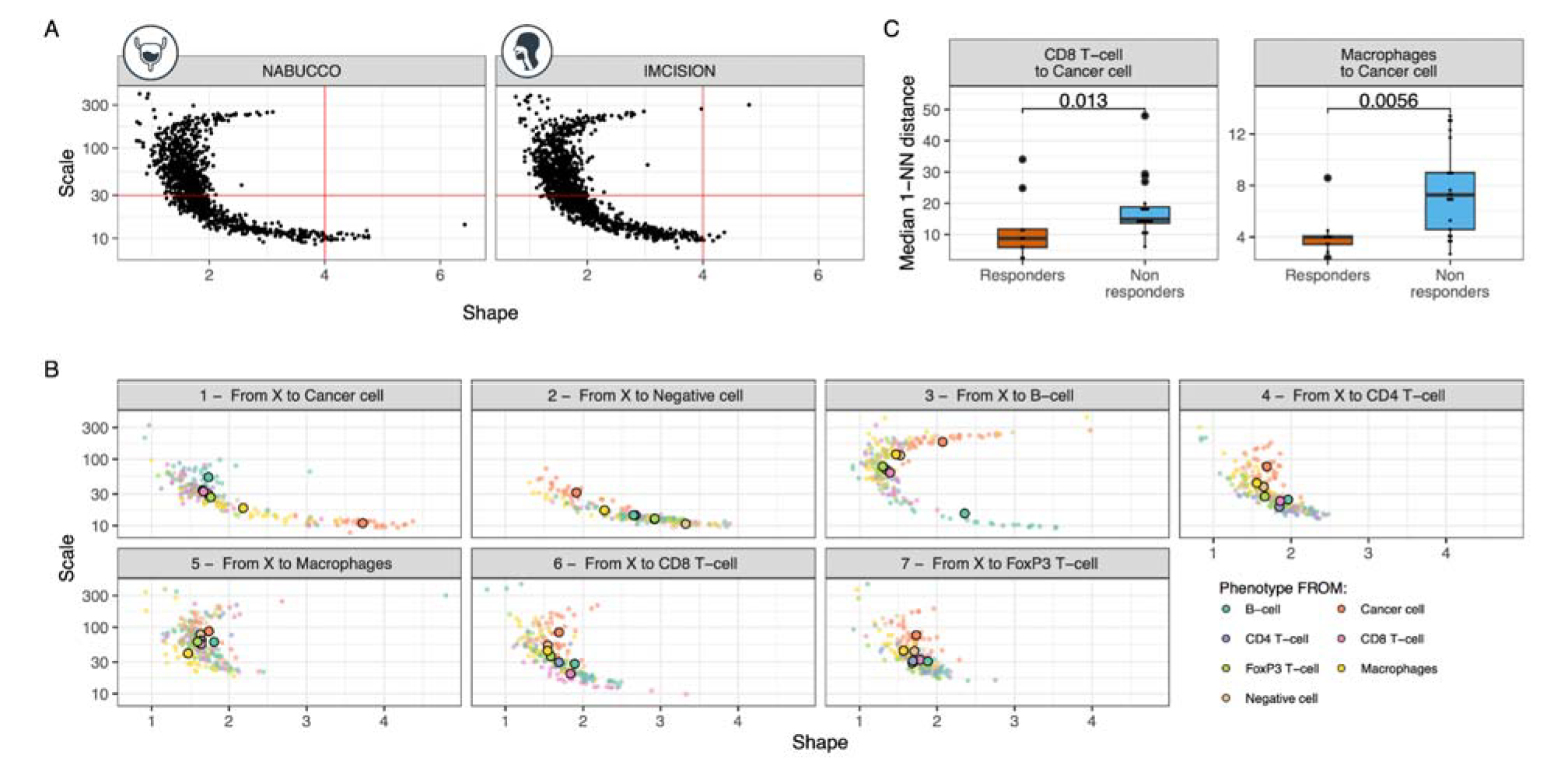
Validation of spatial relationship biomarkers of ICI response in an independent cohort of pre-operative ipilimumab+nivolumab in head and neck cancer. (A) Scale vs. shape SR parameters fitted on the 1-NN distribution for the 25 samples and the 7×7 cell type combinations obtained in the head and neck cancer data (right, IMCISION) and UC data (left, NABUCCO). (B) Scatter plot of the scale-shape parameter space by neighbor cell type (cell to) obtained in the IMCISION trial data. Cohort averages for their associated spatial parameters are highlighted as big dots for each cell type-cell type combination. (C) Median first nearest-neighbor distance distribution in head and neck cancer samples per response group as calculated by the associated shape and scale for the SRs that significantly (FDR<0.04) associated with response in UC and statistical significance by a Mann-Whitney test. Abbreviations: *SR*: spatial relationship; *ICI*: immune checkpoint inhibitors; *1-NN*: first nearest neighbor; *UC:* urothelial cancer.

Next, we evaluated whether the SR biomarkers of ICI response identified in UC were also predictive of reaching a major pathological response upon combination ICI in the HNSCC cohort. The validation was assessed for the strongest biomarkers (FDR<0.04) to maximize the likelihood of validation, which involved the SRs *CD8*^+^ *T-cells to cancer cells* and *Macrophages to cancer cells* from **Figures 4C-D**. Both SRs showed the same direction of association with response in HNSCC (**Figure 5B**) and, importantly, showed a statistically significant association with response after multiple testing correction: CD8^+^ T-cells to cancer cells (FDR_shape_=0.045) and macrophages to cancer cells (FDR_shape_=0.0076, FDR_scale_=0.00094) (**Supplementary Figures 7A-B**), which matches with the spatial proximity behavior (low 1-NN distances) identified in the responding UC tumors (**Figure 4E**). We confirmed the earlier established superiority of the SR metrics over density metrics by showing that, in the HNSCC cohort, immune cell density was not associated with response (**Supplementary Figure 8C**), except for stromal macrophage and CD8^+^ T-cell densities.

In conclusion, the TME spatial biomarkers for pathological response to ICI combination treatment in UC validated in an HNSCC cohort, suggesting that the SRs between CD8^+^ T-cells and macrophages to cancer cells could be an important context-independent biomarker for clinical response to ipilimumab+nivolumab.

## Discussion

Advances in ICI have resulted in pembrolizumab (anti-PD1) becoming the second-line standard of care for advanced UC^3^, and avelumab (anti-PD-L1) as the standard of care for maintenance after chemotherapy treatment^30^. Results from neoadjuvant clinical trials show that patients can have a pathological complete response to only two or three cycles of immunotherapeutic treatment^9, 10, 31, 32^. These promising clinical results need biomarkers that stratify individual patients and improve our understanding of the immunological background of (non-)response. In this study, we provided a comprehensive quantitative exploration of the, thus far, poorly characterized SRs in the UC TME. We show the potential for clinical utility in predicting the response to ICI and provide a quantitative basis for follow-up research.

We found an association between the proximity of the SR from CD8^+^ T-cells to cancer cells and response in UC and confirmed that this relation also holds in HNSCC. In contrast, no differences between response groups in CD8^+^ T-cell density were found, revealing that abundance alone is likely insufficient to explain treatment response. Furthermore, tumors with an immune excluded phenotype exhibit an enrichment of CD8^+^ T-cells at the stroma due to mechanisms preventing T-cells from reaching the tumor. Quantifying immune phenotypes is not trivial due to distinct patterns of exclusion and topography^17, 33^. Our cohort used exclusion ratios and densities to estimate immune phenotypes, and we found no difference between response groups. In contrast, our Weibull parameters served as a distance metric that objectively captures proximity differences between CD8^+^ T-cells and cancer cells between response groups. Our observations suggest that therapeutic strategies that enhance CD8^+^ T-cell migration closer to tumor cells may overcome resistance to ICI. These results align with the observation that immunosuppressive mechanisms, such as TGF-beta signaling, are associated in UC^10, 12^ with a CD8^+^ T-cell excluded phenotype and resistance to ICI^34^. Similar results have been reported in the ICI context for melanoma, in which responding tumors to different ICI treatments were characterized using a 1-NN statistic by proximity between proliferating antigen-experienced CD8^+^ T-cells (CD45RO^+^Ki67^+^) to their closest cancer cell^22^. Moreover, in gynecological and non-small cell lung cancer, the SR between tumor-infiltrating lymphocytes (TIL) and non-TILs (e.g., cancer cells) demonstrated its utility for clinical outcome prediction in an ICI cohort^23^, which is compatible with our observations in UC and HNSCC.

We found that the proximity of macrophages to cancer cells was positively associated with response in the UC and HNSCC cohorts. Interpretation of this candidate biomarker warrants further investigation due to the plasticity and potential pro-or anti-tumorigenic behavior of macrophages^35^, which results in macrophage subtype heterogeneity not covered by our mIF antibody panel. Literature in the ICI context suggests that macrophages can express PD-L1 and PD-1^36^ but can also prevent T-cells from reaching cancer cells^37^. Data from pancreatic cancer suggest that anti-tumorigenic macrophages (M1-macrophages) are closer to cancer cells than pro-tumorigenic macrophages (M2 macrophages)^38^, which indicates that our proximity signal between macrophages and cancer cells in responding tumors may originate from an M1-type macrophage lineage. In locally advanced esophageal squamous cell carcinoma treated with chemoradiotherapy and SHR-1210 (anti-PD-1 ICI), a prognostic signal using the 1-NN statistic median reported PD-L1^+^ tumor cells closer to PD-L1^-^ macrophages associated with a better OS after treatment^24^. Lastly, non-responding tumors were associated with close proximity between B-cells surrounding CD8^+^ T-cells, which is in line with the high baseline expression of genes involved in B-cell signaling we found in non-responding UC tumors in NABUCCO^10^.

We compared the spatial and density metrics’ predictive power to corroborate the SR metrics’ importance. Our results show a superior predictive power for SR metrics and enhance the limited view that count-derived data, such as density or exclusion ratios, provide of the TME. Furthermore, we compared the SR quantifications on our Weibull parameters with the conventional G-function. While both are based on the 1-NN distance statistic, we showed that the G-function dependence on a distance threshold (T) reduced its utility for group comparisons because the associated G-function’s range of values, variance, and predictive power was threshold dependent. Besides, due to the heterogeneity in the G-function evaluations across cell-cell pairwise relationships, there is no unique optimal threshold that maximizes differences between clinical groups of interest for all SRs. Therefore, optimization methodologies for the threshold of choice depend on the SR and cohort, potentially leading to under-or over-fitting and generalization issues. Earlier work on the G-function metric usage for pancreatic cancer grade prediction reported that a single threshold evaluation cannot model all the inherent signals from the data^25^. A higher predictive power could only be achieved by discretizing the G-function at multiple thresholds, which limits its interpretability and utility because of an increased number of summary parameters^25^. On the other hand, our Weibull parameters (shape and scale) allowed for an invariant summary of the SRs without any threshold, which was achieved by, instead of having an empirical summary or discretizing it, modeling all its inherent structure using a curve-fitting approach. Furthermore, the mixed model methodology allowed us to smooth the data and model the parameter variance at a cohort level, making it more suitable for group comparisons when correlating them with clinical phenotypes of interest because of the reduced leverage of outlier samples^39^.

The validation of our TME candidate biomarkers for ICI response identified in UC indicates biologically relevant SR differences consistent across cancer types. Crucially, despite HNSCC being a different organ and morphologically distinct tumor type, showing variability within their biopsy locations (oral cavity, oropharynx), using our proposed spatial approach, we observed similar average SR distances in both tumor types. A combination of pathological complete response and near-complete response defined response to ICI for exploratory analyses in IMCISION (HNSCC). However, in NABUCCO (UC), treatment response was defined as a pathological complete downstaging at the time of surgery. The response rate in IMCISION was lower than in NABUCCO (36% vs. 58%, p=0.04), thus decreasing the statistical power to quantify differences between response groups. Despite these differences and the relatively small sample sizes (24 and 25 for NABUCCO and IMCISION, respectively), the translatability of our findings on the associations between the SR parameters identified in our UC cohort and treatment response in the HNSCC cohort is promising.

Limitations of our study include the number of antibody markers profiled in mIF data, which restricted the types of cells we could detect. Transurethral resections provide a superficial spatial sampling of the whole TME architecture, therefore allowing for a limited profiling of the tumor margin, which is known to contain a higher abundance of immune cells in UC^33^ compared to intratumoral tissue. Nevertheless, the literature suggests that transurethral resection (TUR) material in UC is representative of the whole UC tumor spatial heterogeneity in ∼58-73% of cases at an immune cell density level^40^, but their associated SRs remain yet unexplored. Limitations to our methodological framework include quantifying SRs by studying only the first nearest neighbor and not beyond. Network or graph-based approaches would allow for a broader spatial representation of the TME. However, these topology-based methods usually ignore distances and require more complex SR representations. Furthermore, combinations of samples and pairwise cell type SRs involving noisy distance distributions, such as SRs derived from lowly-populated cells (e.g., FoxP3^+^ T-cells in a subset of UC samples), are excluded from the analysis only when convergence is not reached in the mixed model fitting. However, only 2% of our SRs (24 out of 1176 SRs) were rejected for this reason. While this might have consequences in associations with clinical outcomes of interest (e.g., clinical response), such rare cell types SRs lack robustness. Lastly, our sample sizes are relatively small, and our results warrant further validation in independent and larger cohorts.

In short, our study provides a systematic framework to quantify SRs. It demonstrates that SRs provide a complementary summary of the TME outperforming count-derived metrics, such as density, for identifying biomarkers with a clinical utility. Our results reveal proximity between CD8^+^ T-cells to cancer cells and macrophages to cancer cells as candidate biomarkers for response to neoadjuvant combination ICIs, which have been thus far unexplored and provide a complementary view of the TME that warrants further investigation.

## Supplementary Figures

**Supplementary Figure 1.**
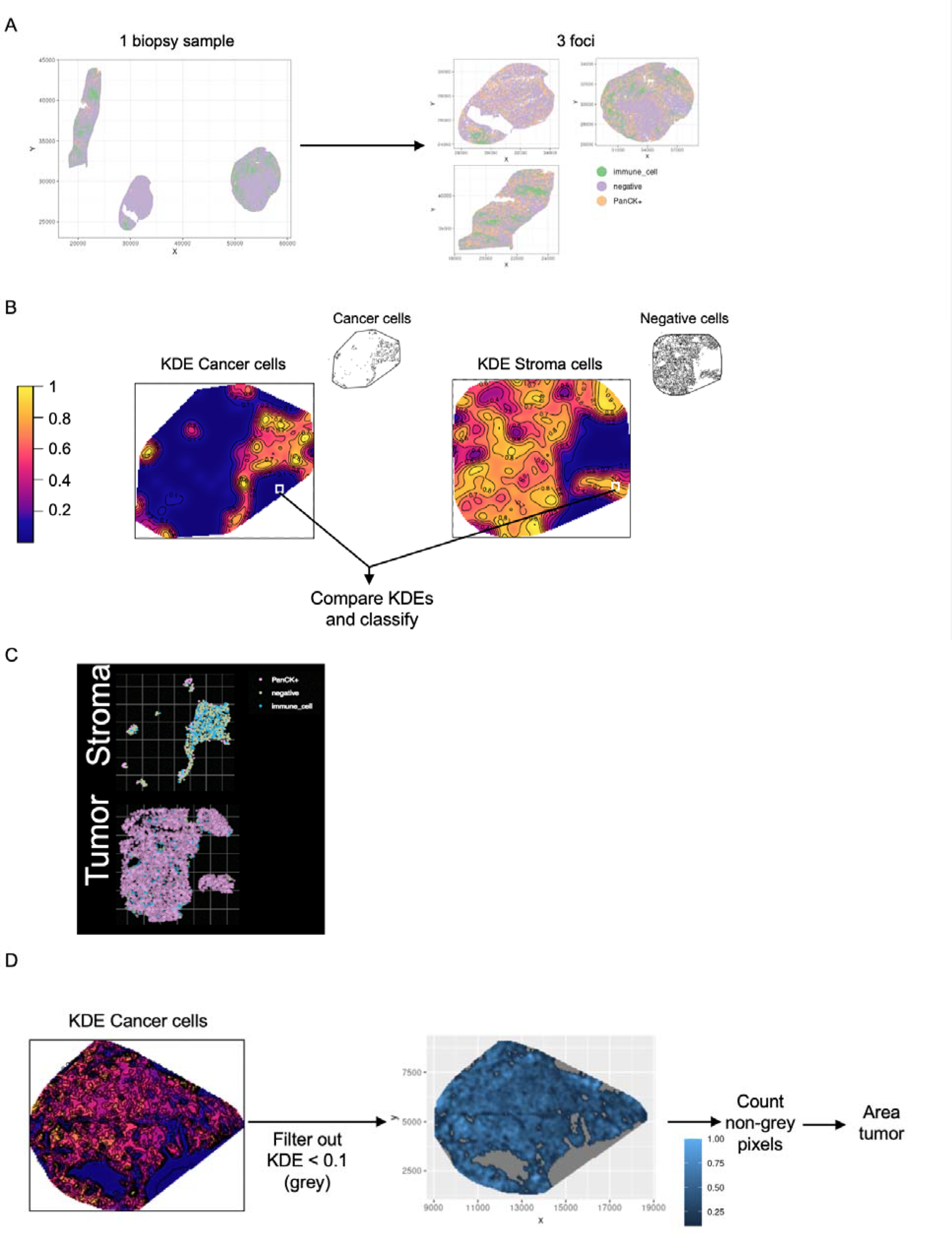
Segmentation of tumor and stroma compartments and tissue area assessment. (A) Visual representation of a biopsy tissue slide (left) that contains multiple tissue islands. Each tissue island was identified using the dbscan algorithm and named foci (right). (B) Tumor and stroma regions segmentation by comparison of the Kernel density estimation (KDE) computed using only cancer cells (left) or stroma cells (right). Prior to segmentation, KDEs were normalized by the maximum value. (C) Example of tumor and stroma classifications. Cells in the top and bottom panels are classified as being in the stroma or the tumor compartment, respectively. Coloring denotes the cell type classification by HALO. (D) Visual representation of the area calculation of the tumor region. First, a KDE using tumor cells was computed and normalized by the maximum value. Then, all the pixels with a KDE below 0.1 were filtered out (drawn as gray in the middle panel). Lastly, all the remaining pixels were counted, and used for the area estimation. Abbreviations: *KDE*: kernel density estimation.

**Supplementary Figure 2.**
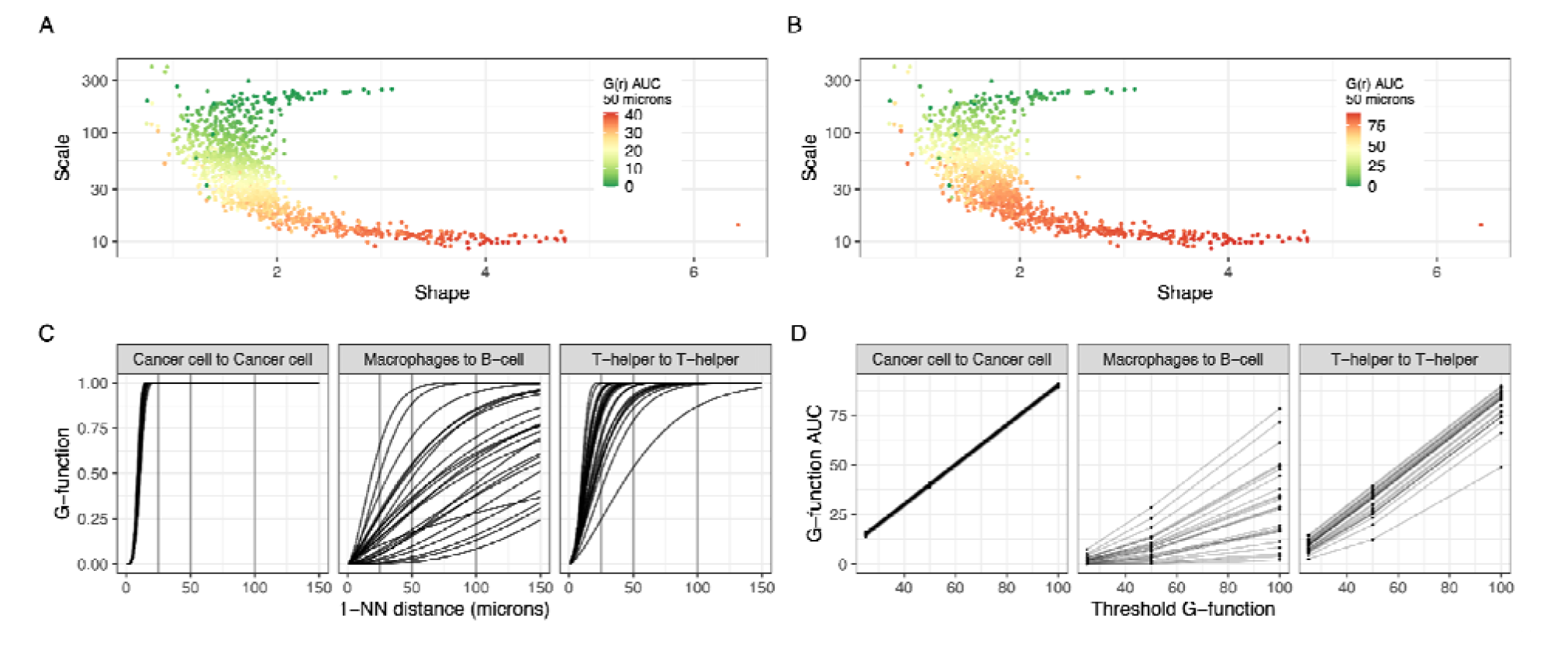
Comparison of spatial parameters derived from the 1-NN distance statistic. (A) Scale vs. shape parameter space associated with the G-function evaluated at 12.5 microns (coloring, G-AUC-12). (B) Scale vs. shape parameter space associated with the G-function evaluated at 50 microns (coloring, G-AUC-50). (C) G-functions for the B-cell to B-cell, CD8^+^ T-cell to Cancer cell, and Macrophages to B-cell SRs. Lines join samples. Vertical lines denote the thresholds 25, 50 and 100 microns. (D) G-AUC-T evaluated at 25, 50 and 100 micron threshold for the examples shown in panel (C). Lines join SRs from the same samples. Abbreviations: *G-AUC-T*: G-function evaluated at a threshold T; *T*: threshold; *AUC*: Area under the curve, *1-NN*: first nearest-neighbor.

**Supplementary Figure 3.**
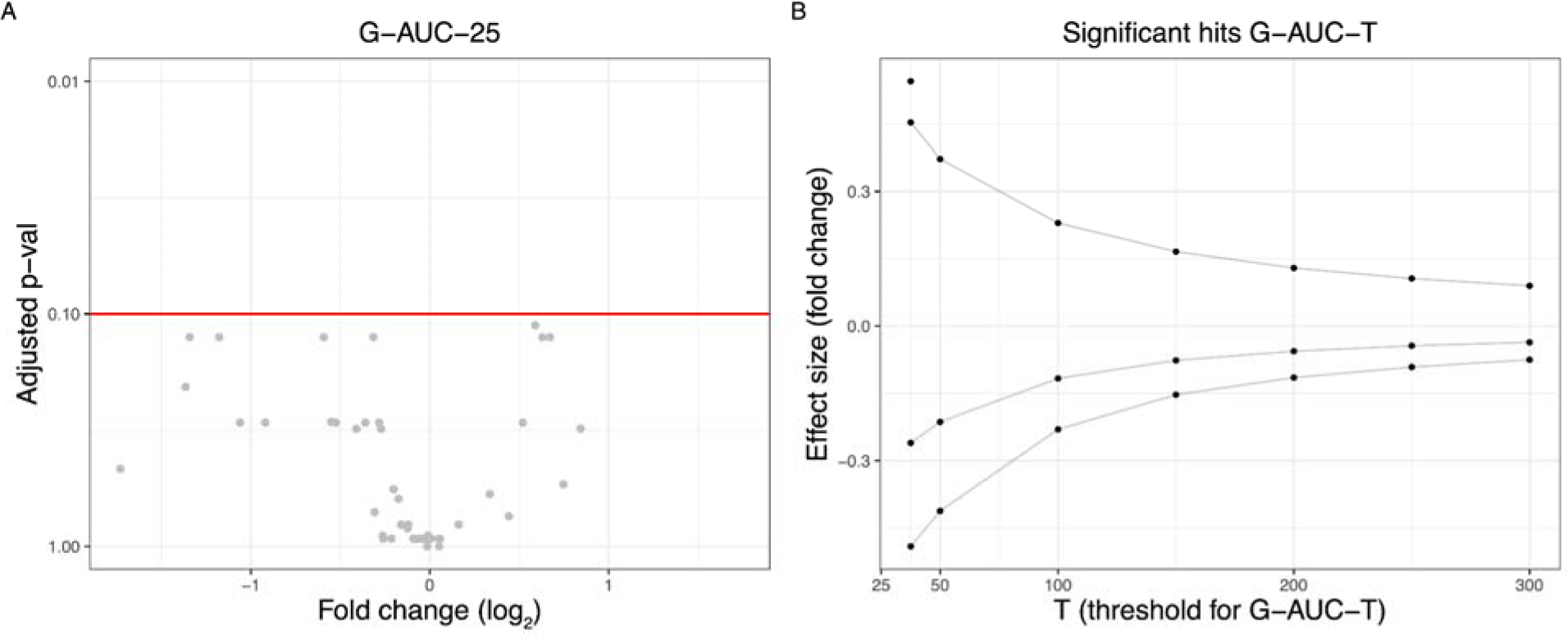
Association of spatial relationships derived from the G-function with response to pre-operative ipilimumab+nivolumab in urothelial cancer. (A) Volcano plot showing the fold change on the G-function parameter evaluated at 25 microns (G-AUC-25) between response groups (x-axis) and statistical significance by Wilcoxon test adjusted by multiple hypothesis testing (y-axis). (B) Fold change (y-axis) on the G-function parameter evaluated at different thresholds (x-axis) between response groups. Only fold changes of significant associations obtained with a Wilcoxon test adjusted by multiple hypothesis testing are shown. Unless otherwise stated, all statistical tests were two-sided. Abbreviations: *G-AUC-T*: G-function evaluated at a threshold T; *T*: threshold; *AUC*: Area under the curve.

**Supplementary Figure 4.**
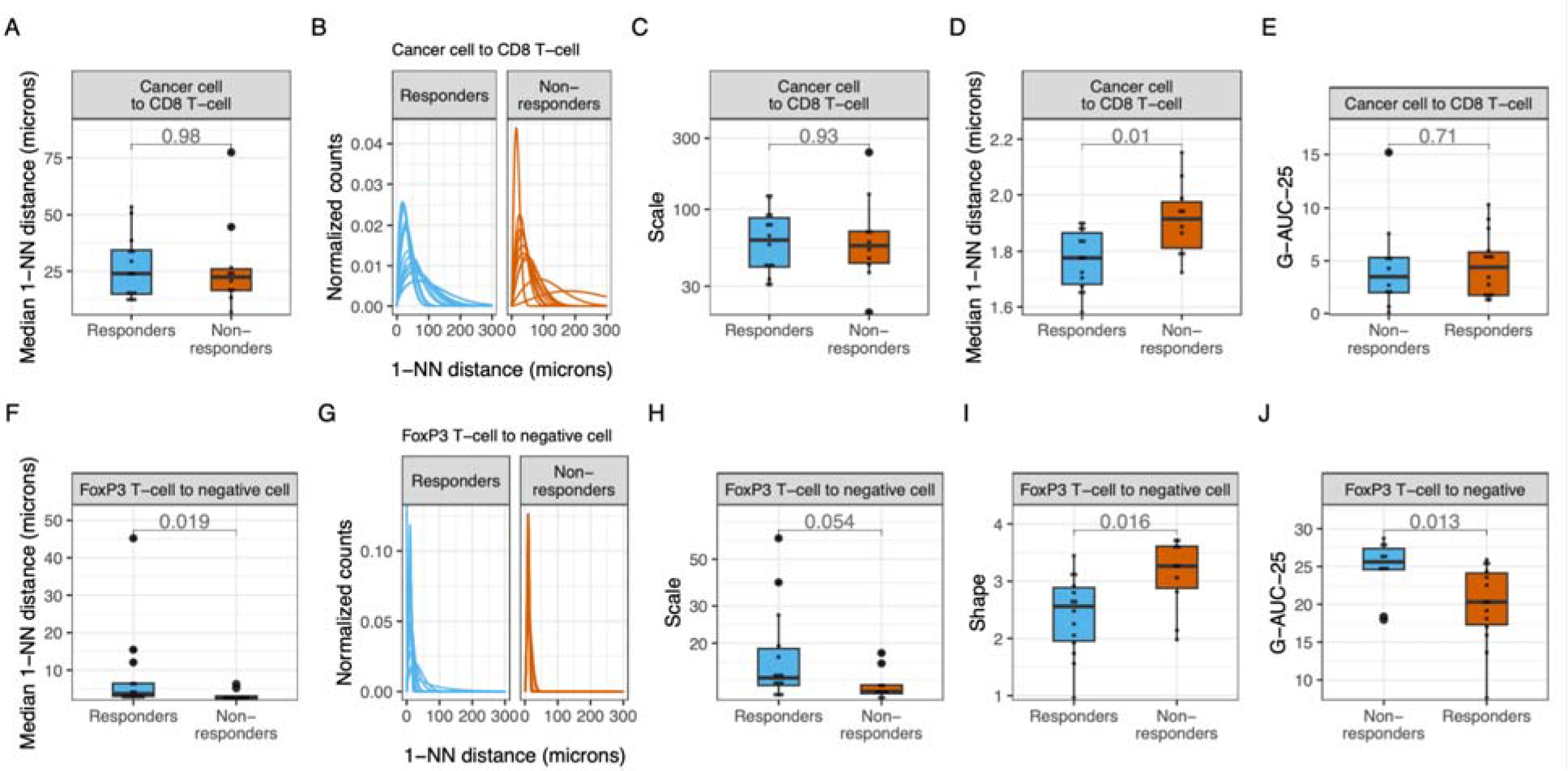
Association of spatial relationships with response to pre-operative ipilimumab+nivolumab in urothelial cancer. (A) Median 1-NN distances, (B) associated 1-NN curves, (C) scale parameters and (D) shape parameter and (D) G-AUC-25 parameters stratified by response groups for the SR from cancer cell to CD8^+^ T-cells (top row) and from FoxP3 T-cells to negative cells (bottom row). Statistical significance was assessed by a Mann-Whitney test (panels A, E, F, J) and a t-test (C, D, H, I). Unless otherwise stated, all statistical tests were two-sided and no adjustments for multiple hypotheses were made. Adjusted p-values: FDR_scale_(cancer cell to CD8^+^ T-cell)=0.98, FDR_shape_(cancer cell to CD8^+^ T-cell)=0.095, FDR_G-AUC-25_(cancer cell to CD8^+^ T-cell)=0.93, FDR_scale_(FoxP3 T-cell to negative cell)=0.21, FDR_shape_(FoxP3 T-cell to negative cell)=0.11, FDR_G-AUC-25_(FoxP3 T-cell to negative cell)=0.13, FDR_G-AUC-25_(FoxP3 T-cell to negative cell)=0.04. Abbreviations: *1-NN*: first nearest neighbor; *SR:* spatial relationship; *G-AUC-25*: AUC of the G-function evaluated at 25 microns; *AUC*: area under the curve..

**Supplementary Figure 5.**
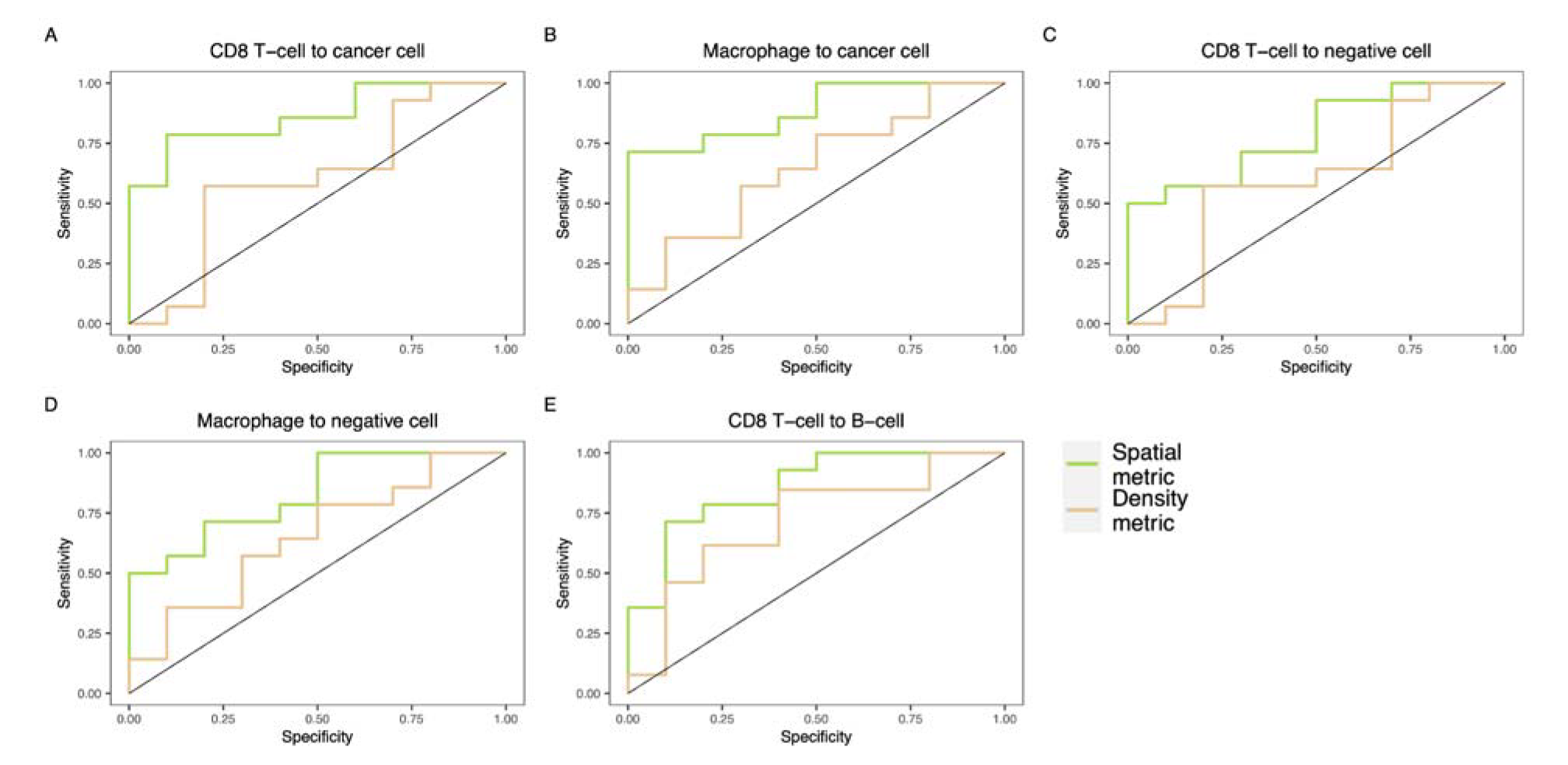
Comparison of the predictive power of associations with response to pre-operative ipilimumab+nivolumab in urothelial cancer between spatial relationships and density parameters. ROC curves of the discriminative power of the spatial and density-related parameters involving significant SRs that are associated with response. ROC-plots from ‘Spatial metrics’ (green) were built upon the predictions of a logistic regression model trained on the shape and scale parameters of the associated SR. ROC-plots from ‘Density metrics’ (orange) were built upon the predictions of a logistic regression model trained on the intratumoral density, stromal density, and exclusion ratio (ratio between stromal and intratumoral density) for the immune cells associated in the SR. Abbreviations: *SR*: spatial relationship; *ROC*: receiver operating characteristic.

**Supplementary Figure 6.**
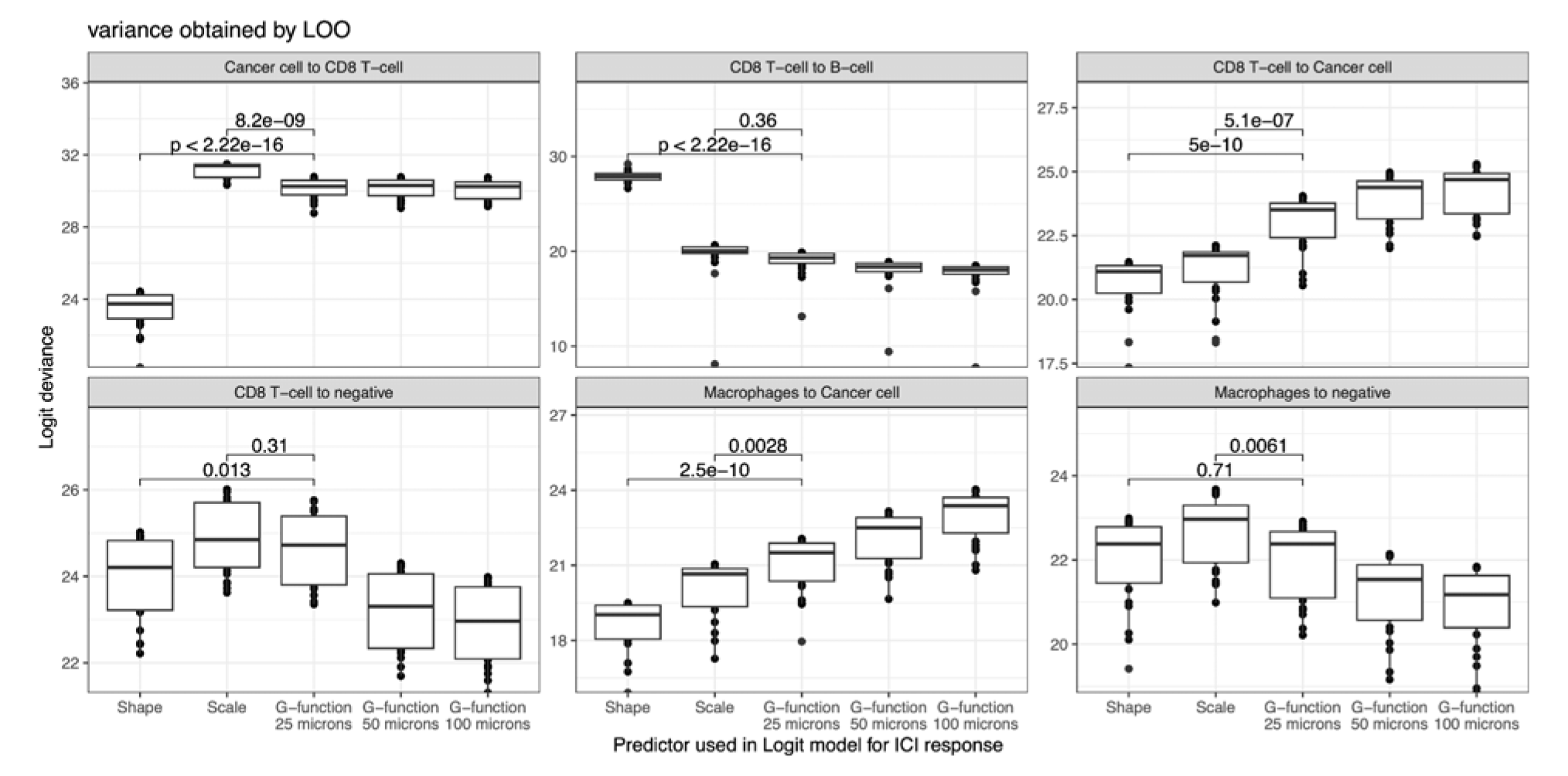
Comparison of predictive models of response to pre-operative ipilimumab+nivolumab in urothelial cancer trained using spatial parameters derived from the 1-NN distance statistic. Logistic regression deviance of a univariate logistic regression model predicting ICI response using as a predictor the shape, the scale, or the G-function evaluated at different thresholds (G-AUC-T). Variability on the AIC was evaluated by leave-one-out cross-validation. Each panel denotes models trained using SRs studied for different cell-cell pairwise relationships. Statistical significance was assessed by a two-sided t-test. Abbreviations: *SR*: spatial relationship; *ICI*: Immune checkpoint inhibitors; *G-AUC-T*: G-function evaluated at a threshold T; *T*: threshold; logit: logistic regression.

**Supplementary Figure 7.**
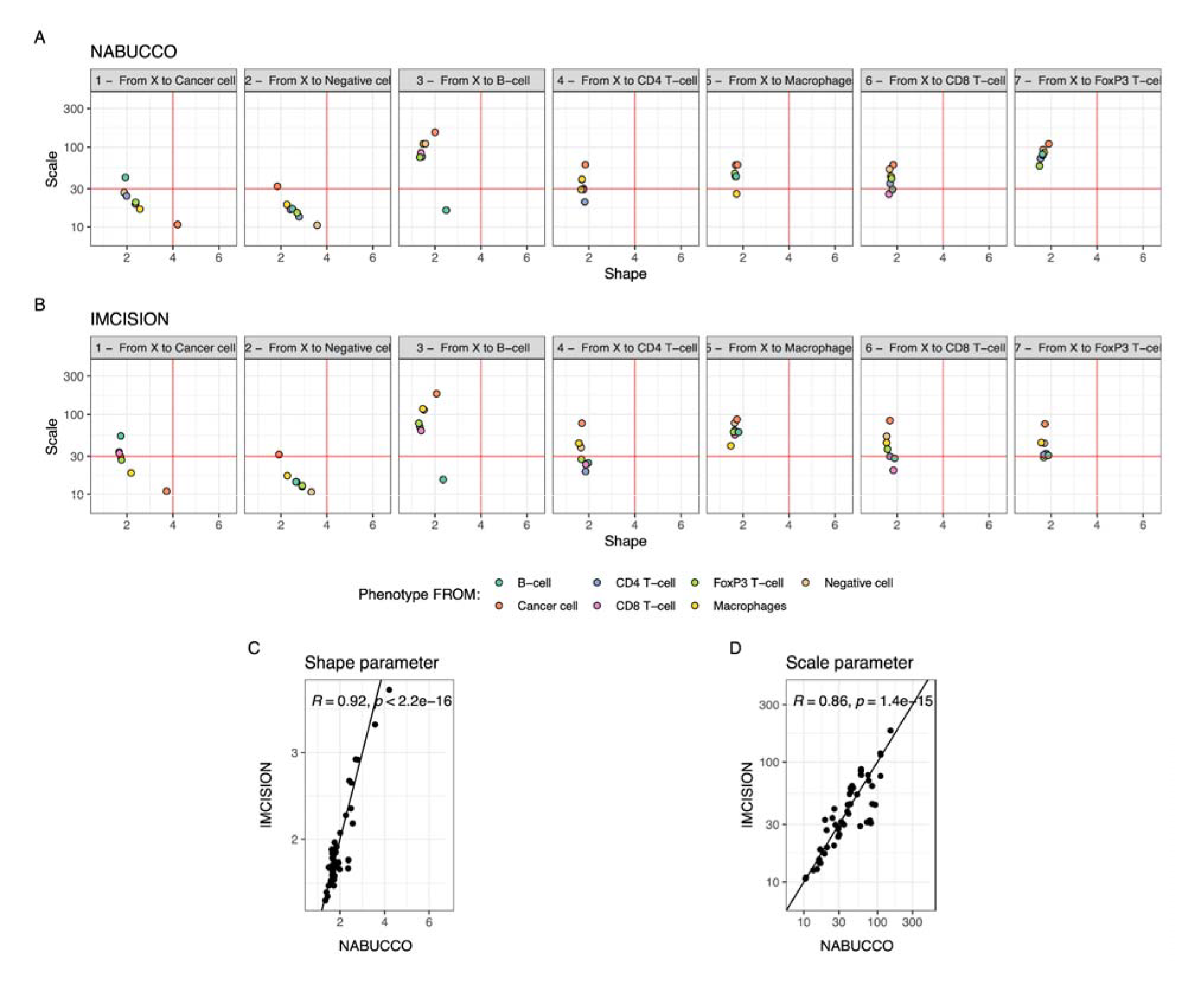
Comparison of spatial relationship parameters derived from pre-operative urothelial cancer (NABUCCO) and head and neck cancer (IMCISION) samples. (A, B) Cohort averages for each SR’s cell type pairwise relationships shape and scale parameters in UC (top, NABUCCO, A) and HNSCC (bottom, IMCISION, B). Each facet represents SRs to a specific target cell type. For instance, the first facet represents SRs studied from any reference cell type to cancer cells, and the color indicates the cell type from which the SR was studied (reference cell type). (C) Correlation between shape parameters quantified in NABUCCO (x-axis) and IMCISION (y-axis). Pearson’s coefficient and correlation p-value are shown in the plot. (D) Correlation between scale parameters quantified in NABUCCO (x-axis) and IMCISION (y-axis). Pearson’s coefficient and correlation p-value are shown in the plot. All statistical tests were two-sided. No adjustments were made to correct for multiple comparisons. Abbreviations: *SR*: spatial relationship; *UC:* urothelial cancer; *HNSCC*: head and neck squamous cell carcinoma.

**Supplementary Figure 8.**
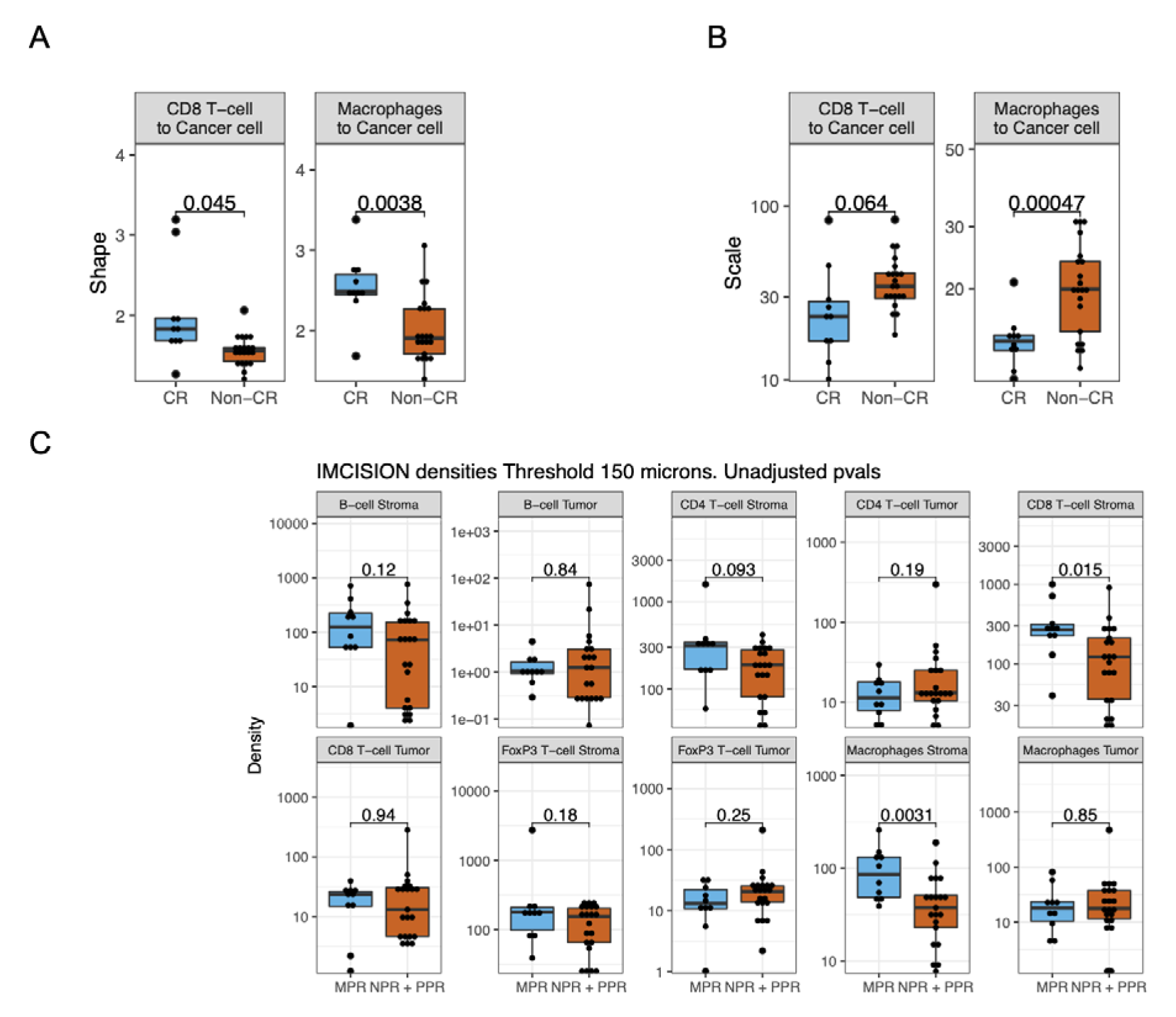
Validation of SR biomarkers of ICI response in head and neck cancer cohort (IMCISION trial). (A) Shape parameters for the SRs from CD8^+^ T-cells to Cancer cells (left) and from Macrophages to cancer cells (right) between ICI response groups. Adjusted p-values: FDR^CD8^ ^T-cell^ ^to^ ^Cancer^ ^cell^=0.045, FDR^Macrophages^ ^to^ ^Cancer^ ^cell^=0.0076. (B) Scale parameters for the SRs from CD8^+^ T-cells to Cancer cells (left) and from Macrophages to cancer cells (right) between ICI response groups. A two-sided t-test was used for comparisons between response groups. Adjusted p-values: FDR^CD8 T-cell to Cancer cell^=0.064, FDR^Macrophages to Cancer cell^=0.00094. (C) Immune cell densities between response groups in IMCISION. A two-sided t-test was used for comparisons between response groups. Unless otherwise stated, no adjustments were made for multiple hypothesis testing. Abbreviations: SR: spatial relationship; *ICI*: immune checkpoint inhibitors.

## Supplementary Tables

### Supplementary Table 1

Density parameters and exclusion ratios quantified from the NABUCCO cohort.

Abbreviations: CR: Responder; NCR: Non-responder.

**File: STable1 NABUCCO_density_parameters.xlsx**

### Supplementary Table 2

Spatial relationship parameters quantified from the NABUCCO cohort (shape, scale, and G-function evaluated at 25, 50 and 100 microns).

Abbreviations: CR: Responder; NCR: Non-responder; AUC: area under the curve; SR: spatial relationship.

**File: STable2 NABUCCO_spatial_parameters_with_G_function.xlsx**

### Supplementary Table 3

Spatial relationship parameters quantified from the IMCISION cohort.

Abbreviations: CR: Responder; NCR: Non-responder; SR: spatial relationship.

**File: STable3 IMCISION_spatial_parameters.xlsx**

## Acknowledgments

We thank the facilities and staff at the Netherlands Cancer Institute (NKI) that contributed to this project, including the Core Facility Molecular Pathology and Biobanking (I. Hofland, S. Vonk, and W. Kievit), the Pathology department (I. M. Seignette, C. van Rooijen, M. Almekinders, L. Smit, E. Hooijberg, and J. Sanders), the Urology department (K. Hendricksen, H. van der Poel) and the Research High-Performance Computing facility. We thank all patients who participated in the NABUCCO and IMCISION clinical trials and the study teams involved. This research at the NKI was supported by an institutional grant from the Dutch Cancer Society (KWF) and the Dutch Ministry of Health, Welfare, and Sport.

## Data availability

Multiplex immunofluorescence primary data used for this manuscript will be made available upon reasonable academic request and within the limitations of the provided informed consent by the corresponding author upon reasonable request upon acceptance. The institutional review board of the Netherlands Cancer Institute will review each request. After approval, the researcher must sign the Netherlands Cancer Institute data access agreement.

## Code availability

Code to reproduce the main findings will be made available in a Github repository.

## Author contributions

Interpreted and addressed clinical data NvD, JLv, CLZ, BvR, MSvdH

Collected clinical data NvD, JLv, CLZ, MSvdH

Collected data NvD, MLvM, DP, EH, AB

Performed bioinformatic and statistical analysis AG-J, YL, DJV

Performed HALO image analysis NvD

Interpreted data AG-J, NvD, DJV, MSvdH, LW

Administrative, technical, or material support DP, EH

Performed multiplex immunofluorescence experiment DP

Performed HALO image analysis NvD

Supervised multiplex Immunofluorescence experiments and assessed tissue availability EH, AB, MLvM

Obtaining funding MDvdH

Supervision DJV, MSvdH, LFAW

Wrote the manuscript together with all co-authors who approved data accuracy AG-J, DJV, MSvdH, LFAW

Critical review of the manuscript and scientific input All authors

## Materials and Methods

### Urothelial cancer study population and treatment (NABUCCO trial)

Twenty-four pre-treatment Urothelial Cancer (UC) tumor samples from the NABUCCO trial (NCT03387761, Cohort 1) were used for analyses. In the trial patients underwent a combination treatment (2 or 3 cycles) of ipilimumab (anti-CTLA-4) and nivolumab (anti-PD-1) prior to surgical resection. The trial cohort consisted of high-grade stage III muscle-invasive urothelial cancer (cT3-4aN0M0 or cT1-4aN1-3M0). Details of the trial are reported^10^.

Response to treatment was evaluated by pathological response assessment on radical surgery. Tumors with a complete pathological response (ypT0N0) or residual disease (<=ypT1N0) were classified as responders (n=14), and tumors with a >=ypT2N0 were classified as non-responders (n=10).

### Head and Neck squamous cell carcinoma population and treatment (IMCISION trial)

Thirty-one head and neck squamous cell carcinoma (HNSCC) tumor samples of multiple subsites (oral cavity n=27, oropharynx, n=4) were obtained from the IMCISION trial (NCT03003637). Patients underwent either two cycles of nivolumab (Arm A, n=6) or a combination treatment of 2 cycles of ipilimumab (anti-CTLA-4) and nivolumab (anti-PD-1) (Arm B, n=25) prior to surgical resection. The trial cohort consisted of HNSCC tumors with a histological grade T2LT4N0LN3b and metastatic grade M0 primary or recurrent of mostly HPV-negative head and neck squamous cell carcinoma (HPV negative, n=23; HPV positive, n=2). Details of the trial can be found elsewhere^27^. Only samples from Arm B (n=25) were analyzed in this manuscript.

Response to treatment was evaluated by pathological response assessment on surgery and by comparison of tumor cells decrease from baseline to on-treatment samples^27^. Tumors with <=10% tumor cell percentage (TCP) at surgery and a decrease of 90-100% in tumor cells from baseline to on-treatment were classified as major pathological responders (MPR, n=9); tumors with <=50% TCP at surgery and a decrease of 50-89% in tumor cells from baseline to on-treatment were classified as partial pathological responders (PPR, n=1); else tumors were classified as no pathological responders (NPR, n=15). Patients with an MPR were classified as Responders, and patients with a PPR or NPR were classified as Non-Responders.

### Multiplex Immunofluorescence

Multiplex immunofluorescence (mIF) was performed on pre-treatment formalin-fixed paraffin-embedded (FFPE) sections and assessed on an immune panel (DAPI, PanCK, CD8, CD3, FoxP3, CD20, CD68) as previously described for UC^10^ (NABUCCO) and HNSCC^27^ (IMCISION). The experimental protocol and data processing is reported elsewhere^10, 27^. Upon mIF profiling, cells were segmented by marker positivity:

| Marker positivity* | Cell type |
| --- | --- |
| PanCK<sup>+</sup> | Cancer cell |
| CD3<sup>+</sup>CD8<sup>+</sup> | CD8 T-cell |
| CD3<sup>+</sup>FoxP3<sup>+</sup> | FoxP3 T-cell |
| CD3<sup>+</sup>CD8<sup>-</sup>FoxP3<sup>-</sup> | CD4 T-cell |
| CD68<sup>+</sup> | Macrophage |
| CD20<sup>+</sup> | B-cell |
| Negative for all markers above | Negative cell |

\*all cells were DAPI positive

Multiplex immunofluorescence in IMCISION was assessed as in NABUCCO and the experimental protocol is published in the original manuscript^27^. To ensure consistency in the mIF spatial data between the HNSCC and UC cohorts, including similar tumor purity, for each sample from IMCISION we aligned analysis methods by discarding stromal tissue residing beyond 150 microns distance from the tumor tissue by filtering out all cells classified as belonging to the ‘Stroma compartment’ (by segmentation) if their closest cancer cell lay beyond 150 microns by using the *nncross* method from *spatstat*.

The data subjected to downstream analysis represented the position of each cell in the tissue (x-and y-coordinates of the nuclei) and its corresponding cell type.

### Segmentation of tumor and stroma compartments

To segment the tumor and stroma regions from each tissue, we first split each individual “tissue island” from each sample biopsy using dbscan v1.1-6 (density-based spatial clustering of applications with noise) by setting the size of the epsilon neighborhood to 300 and the minimum number of points in the epsilon neighborhood to 50 (**Supplementary Figure 1A**). Each tissue island was named foci.

To segment the tumor and stroma compartments for each focus, we first computed the kernel density estimation (KDE) of the point pattern defined by cancer cells (KDE_tumor_), and by negative cells (KDE_negative_). The KDE was estimated as implemented by *density* in the stats v3.6.3 package. The smoothing bandwidth for the KDE was optimized using likelihood cross-validation as implemented in *bw.ppl* in spatstat v1.64-1. Then, the KDEs were normalized to their maximum value (KDE_tumor=_ KDE_tumor_ / max(KDE_tumor_)) to allow comparison between the KDE of the tumor and negative cells. To segment the tissue, for each position populated by a cell, we compared both KDEs, and classified them as “Tumor” when KDE_tumor_ > KDE_negative_ and as “Stroma” otherwise (**Supplementary Figure 1B-C**).

### Calculation of tumor and stroma compartment areas

To compute the covered area by each segmented tissue compartment (“tumor” or “stroma”), we first computed the kernel density estimation (KDE) of the point pattern defined by the cancer cells (KDE_tumor_) and normalized by maximum KDE intensity. Then, to compute the area of the tumor compartment, we filtered out all the KDE pixels with a normalized intensity < 0.1. We selected this threshold based on visual exploration for all cells. We then estimated the tumor compartment area as the aggregated area of all non-filtered pixels from KDE_tumor_ (thus with intensity >= 0.1).

To compute the total tissue area, we also computed the KDE_negative_ and normalized it by the maximum value. We then summed the KDE_tumor_ and KDE_negative_, and filtered out pixels with a normalized intensity < 0.1. We then estimated the Total tissue area as the aggregated area of non-filtered pixels from (KDE_tumor_ + KDE_negative_) (intensity >= 0.1).

To estimate the area of the “stroma” compartment, we subtracted the “tumor area” from the “total tissue” area (**Supplementary Figure 1D**). This process was performed for each foci.

### Spatial analysis: quantification of the first nearest neighbor (1-NN) distance distribution

The spatial relationships between all cells within the tumor microenvironment were studied using the first-nearest neighbor (1-NN) statistic as implemented in *spatstats*. In brief, the approach is studied from a reference cell type to a neighbor cell type. For each reference cell type (“*cell type from”*), the distance to the closest neighbor cell type (“*cell type to”*) was measured using *nndist* (**Figure 1E**). Then, we constructed a histogram from the vector of 1-NN distances. We smoothed the distribution by sliding a 5-micron window across the 1-NN histogram and iteratively counting the frequency of the 1-NN distances for each micron. We normalized the distribution to achieve a unit area under the curve (AUC) by dividing for the numerical AUC. SRs were quantified using the data from the whole tissue slide (i.e., not making a distinction beween tumor and stroma compartments).

### Spatial analysis: fitting of Weibull distribution to the 1-NN distances vector

To summarize the 1-NN distance distribution, we fitted a Weibull distribution to the empirical probability density function (PDF), which is a 2-parameter distribution based on (the positive parameters) shape and scale, defined as:

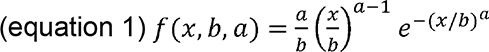

Here, “*b”* denotes the scale, and “*a”* denotes the shape.

We implemented a methodology based on a functional data analysis approach to fit the Weibull distribution. First, to have an initial estimate of the distribution parameters for each patient (n=24) and cell type-cell type combination (n=49), we employed maximum likelihood estimation (MLE) using *fitdist* as implemented on fitdistrplus v1.1.3 package to have an initial estimate of the scale and shape parameters. Then, for each pairwise cell type relationship (cell type from vs. cell to), we implemented a non-linear mixed effect model (nlme v3.1-144) to fit a Weibull distribution on all patient samples, having the shape / scale intercept as fixed effects (*fixed = a+b∼1*) and allowing a random effect for the scale/shape on each sample (*random = list{sample=pdDiag(a+b∼1)}*) by modeling the correlation structure of the random effects a with diagonal positive-definite matrix. The *nlme* model was implemented by maximizing the restricted log-likelihood (method=’REML’), with the parameters set to their default values and the control values set as:

- 1000 maximum iterations for the optimization algorithm (maxIter = 1000).
- 200 maximum iterations for the optimization step which is inside the nlme optimization process (msMaxIter=200).
- 1e-1 tolerance for the PNLS step convergence criterion (pnlsTol=1e-1).
- 1e-6 tolerance for the convergence criterion in nlme algorithm (tolerance=1e-6).
- Nonlinear minimization optimizer (opt="nlm").

To filter out low-quality distributions that lead to non-convergence of the models, we sequentially filtered out samples based on the number of cells (n) from the reference cell type (*nFROM*) or neighbor cell type (*nTO*). First, we fitted a model with the data for the 24 samples. If convergence was not achieved, we filtered out all samples with fewer than 20 cells (*nFROM* < 20 or *nTO* < 20). If the model still did not converge, we repeated filtering samples with fewer than 50, 70, or 100 cells. The approach allowed us to model as much data as possible unless the goodness of fit was compromised. For 24 spatial relationships for which a data fit could not be carried out (2% of the total combinations of data points from 24 samples and 7×7 cell type pairwise relationships) in our cohort.

Because the positivity constraint for the Weibull distribution parameters (b>0, a>0) could not be optimally implemented using a constrained non-linear mixed effect model, we re-parameterized the shape and scale parameters to force the values to be positive. In brief, we re-parameterized the scale and shape employing new parameters *A* and *B*, which were unconstrained:

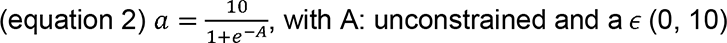

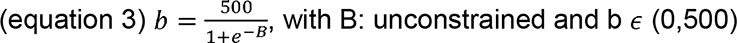

Here, “*a”* is the shape and *“b”* is the scale.

Unless otherwise stated, the SR parameters reported in the manuscript correspond to the ones calculated on the first-nearest neighbor distributions using the spatial distribution of the whole tissue slide (thus, not using data stratified by tumor or stroma).

Sources of the variability of the SR parameters were also quantified, as reported in **Extended Methods**.

### Spatial analysis: Computation of the G-function

Alternatively to fitting a distribution to the 1-NN distance distribution, we computed the G-function, which is defined as the cumulative distribution function (CDF) of the first-nearest neighbor distance distribution:

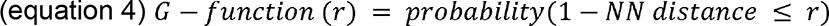

The G-function was computed as implemented by *gest* in the *spatstat* package. Because our SRs were studied between different cell types, we used the multitype nearest-neighbor function G_ij_(r) as implemented in *Gcross*, which was calculated from the first nearest neighbor distances from a cell of type i to the nearest point of type j. Then, to summarize the G-function, we computed the Area Under the Curve (AUC) of the G-function, named G-AUC-T, at different thresholds T’s, which included 25, 50, and 100 microns (i.e., G-AUC-25, G-AUC-50, and G-AUC-100, respectively).

### Spatial analysis: Weibull-derived G-function

Using the properties of the Weibull distribution, we analytically constructed the Cumulative Distribution Function (CDF) using the shape and the scale parameters:

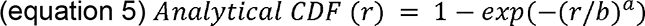

Here, *b* and *a* denote the scale and shape, respectively, and *r* denotes the first nearest neighbor distance.

The Analytical CDF is analogous to the Analytical G-function. The AUC of the Analytical CDF at was evaluated at different distance thresholds T’s, and referred to in the manuscript as Weibull-G-AUC-T.

### Comparison of discriminative power between spatial and density-related parameters

We compared the predictive power between ICI response groups of the spatial-related parameters (shape, scale), density-based parameters (intratumoral and stromal immune cell density) and exclusion-based parameters (ratio between stromal and intratumoral immune cell densities). First, for each set of parameters (e.g., spatial-related parameters), we trained a logistic regression model using *glm* to predict response. Then, a ROC curve was built using the logistic regression’s fitted values (probabilities) using pROC v1.17.0.1. For each ROC-AUC, we tested whether the AUC was significantly different from AUC=0.5 using a two-sided Wilcoxon signed-rank test between cases and controls. We used a bootstrapped approach (n=500) to estimate the confidence intervals of the ROC-AUCs and to test whether the ROC-AUCs from the spatial parameters were significantly greater than the ROC-AUCs from either the density or the exclusion ratio parameters.

### Comparison of discriminative power between Weibull-derived or G-function-derived parameters

Logistic regression deviance of Weibull parameters (shape, scale) and G-function parameters (G-AUC-T evaluated at different Ts) was evaluated by training univariate logistic regression (LR) models using data from each feature as a predictor, and clinical response labels as the dependent variable. Logistic regression deviance (logit deviance) was assessed, in which lower values denote better model fits. The uncertainty of the AIC was estimated by performing a leave-one-out (LOO) variant of the analysis. A two-sided student’s t-test tested statistical significance.

### Spatial biomarker validation in HNSCC

We validated our top SR biomarkers associated with clinical response in our UC cohort (NABUCCO). First, we selected the top two biomarkers identified using our pipeline in UC (FDR < 0.04 in either the shape or scale parameter, which yielded the SRs *CD8*^+^ *T-cells to cancer cells* and *Macrophages to cancer cells).* Second, we evaluated the shape and scale parameters for the biomarkers mentioned above between clinical response groups in the external HNSCC (IMCISION) cohort using a two-sided t-test and adjusted for multiple hypothesis testing using the Benjamini-Hochberg method. Then, we evaluated the median 1-NN distances using the analytical derivation from the shape and scale parameters:

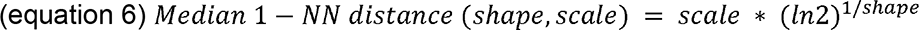

### Statistical analysis

Unless otherwise stated, a two-sided student’s t-test was used for group comparisons. We modeled density and count data in a logarithmic space. For spatial data, the shape of the Weibull parameters (*a*) was modeled on a non-logarithmic scale, and the scale (*b*) was studied on a logarithmic scale. Multiple hypothesis testing corrections were done using the Benjamini-Hochberg method. Unless otherwise noted, statistical significance was defined as p<0.05 and False Discovery Rate (FDR) < 0.10 (10%), and all statistical tests were two-sided. All statistical analyses were performed in R 3.6.3. The following packages were used in this study:

- Spatstat 1.64 ^28^
- Dplyr 1.0.4
- Fitdistrplus 1.1.3
- Patchwork 1.1.1
- Survival 1.3.24
- ComplexHeatmap 2.2
- Circlize 0.4.12
- Glmnet 4.1.1
- RColorBrewer 1.1.2
- Nlme 3.1.144
- Spastat 1.7.0
- Ggpubr 0.4.0
- Ggrepel 0.9.1
- Plyr 1.8.6
- Tidyverse 1.3.0
- Ggplot 2.3.3.3
- Tibble 3.0.6
- ggrastr version 1.0.1
- pROC 1.17.0.1

## Appendix

## Supplementary Notes

### Supplementary Note 1: Quantifying sources of variation of the spatial relationship parameters

To investigate whether SR metrics are susceptible to changes in immune cell density, we performed a simulation study based on the real patient mIF data (Supplementary Methods, *Spatial analysis: a simulation study to quantify spatial relationship parameters sources of variation*). We preserved the spatial positions of cells (but not their identity) and the tumor and stroma delineation. We then placed immune cells at an increasing density at the preserved cell positions for the cohort. **Supplementary Notes Figure 1A** shows a representative sample where the immune cells are only placed in the cancer compartment, i.e., replacing a cancer cell with an immune cell. We varied the percentage of immune cells in the tumor compartment from 1 to 10%, where the percentage is a fraction of the total number of cells on a slide. This range is representative of the variation in immune cell fraction we observed in our cohort (2-1583 immune cells/micron^2^).

We started by simulating multiple intratumoral immune cell densities, which included *Low* (1% of immune cells), *Medium* (4% of immune cells), or *High* (7% of immune cells) densities (**Supplementary Notes Figure 1A**). We then estimated all the pairwise SR parameters between cancer, immune and negative cells. As expected, an increase in immune cell density decreased the 1-NN distances between immune cells (**Supplementary Notes Figure 1B**, and **Supplementary Notes Figure 2A**, right panel). The associated medians for each 1-NN spatial distribution shown in **Supplementary Notes Figure 1B** confirmed these findings (**Supplementary Notes Figure 2A**). In contrast, changes in immune cell density did not affect the 1-NN curves nor the metrics quantifying the SR from immune to cancer cells (**Supplementary Notes Figure 2A**). Our findings showed little variation between the different patient samples (**Supplementary Notes Figures 3A-B**). In summary, we found that density can affect the SR metrics, but that is highly dependent on the cell types involved and the directionality of the relationship.

We can conceive the following immune cells spatially distributions giving rise to the following configurations of immune phenotypes^29^. First, being Excluded (higher immune cell abundance in the stroma), Inflamed (high immune cell abundance in the tumor), and Desert (low immune cell abundance in the tumor and stroma). In order to quantify the effect of immune phenotypes on the SR metrics, we simulated the associated extreme scenarios of the immune phenotypes, and named them as *Excluded* (immune cells only in the stroma and not in the tumor), *Mixed* (immune cells both in the stroma and tumor), and *Inflamed* (immune cells only in the tumor), all at distinct immune cell densities (**Supplementary Notes Figure 1C**). As expected, the different immune phenotypes have a distinct effect on the SR of immune to cancer cells. In contrast, the SR immune to immune cells remains unaffected, similar to the observation when the immune cell density was varied (**Supplementary Notes Figures 1D, 2B**). Notably, these effects were remarkably stable across the complete cohort (small variance in **Supplementary Notes Figure 3A-B**), indicating that the specific positions of cells and the arrangement of tumor and stroma in a particular tumor do not have a strong effect on the simulated SRs. Cases from a particular configuration (e.g., *Inflamed*) consisted of aggregated simulations of distinct immune cell densities compatible with such configuration to assess the true effect between immune phenotypes in our comparisons (e.g., *Inflamed* vs. *Excluded*). Again, little sample variability within the whole cohort was observed between the associations of immune phenotype perturbation and SR parameters (**Supplementary Notes Figure 3C-D**).

Next, we compared the SR metrics for homogeneously and heterogeneously distributed immune cells. Examples of such instances include T-cells homogeneously infiltrating a tumor or B-cells forming immune cell clusters resulting in non-homogeneous distributions of B-cells. We simulated these two scenarios (**Supplementary Notes Figure 1E**) and compared the associated SR metrics. We found that immune cell clustering can affect the SR metrics from immune to immune cells and from immune to cancer cells (**Supplementary Notes Figures 1F**, **2C**, **3E-F**).

To summarize our findings, we aggregated simulations for all the cell type pairwise relationships and samples (**Supplementary Notes Figure 3G**). Changes in the SR parameters upon the posed perturbations highly depend on which reference and target cell types were studied (**Supplementary Notes Figure 4**). The SRs between abundant cell types (i.e., cancer and negative cells) were not altered upon such perturbations. In contrast, the SRs between rare cell types (i.e., immune cells) were affected by the density or local clustering perturbations and SRs from abundant to immune cells were affected by the density perturbations. Moreover, immune phenotypes and clustering perturbations affected SRs from immune to cancer cells. In conclusion, we identified multiple factors that affect the magnitude of SR parameters, which depending on the cell type context being studied, need to be considered in downstream analysis.

## Supplementary Notes Figures

**Supplementary Notes Figure 1.**
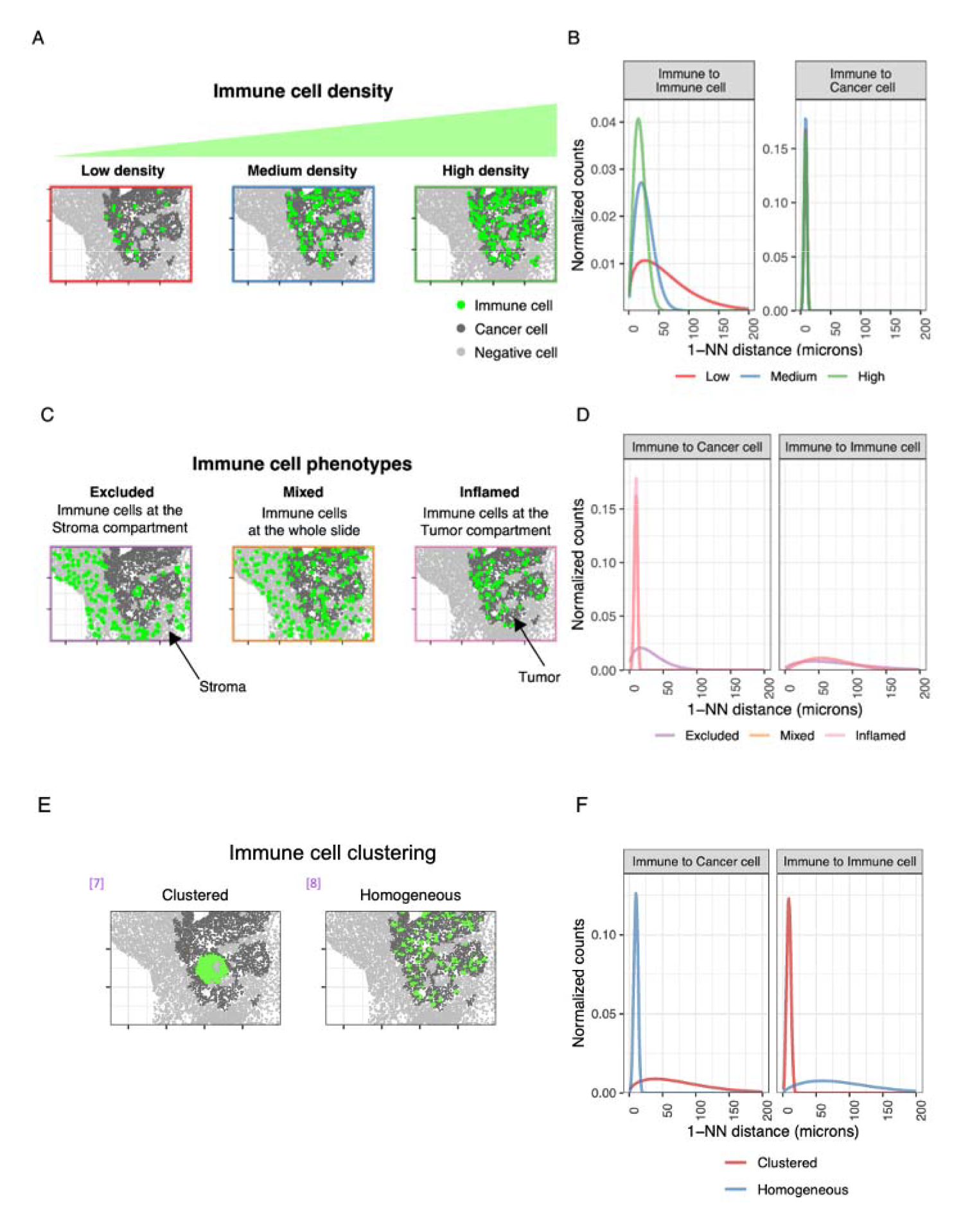
Simulation study to quantify sources of variation of the spatial relationship parameters. Representative data for a sample: (A) Immune cell density was simulated at different values. (B) 1-NN curves vs immune cell density for three representative SR parameters. (C) Immune cells were simulated to be present only in the stroma region (*Excluded*, left), both at the stroma and the tumor region (*Mixed*, middle), and only in the tumor region (*Inflamed*, right). (D) 1-NN curves vs immune cell phenotypes for three representative SR parameters. (E) Immune cell clustering was simulated with values *Clustered* (left) and *Homogeneous* (right) spatial pattern. (F) First-nearest neighbor distance curves for the clustered and homogeneous examples from panel C for the SRs studied from Immune to cancer cells and from Immune to immune cells. Abbreviations: *SR*: spatial relationship; *ICI*: immune checkpoint inhibitors; *1-NN*: first nearest-neighbor.

**Supplementary Notes Figure 2.**
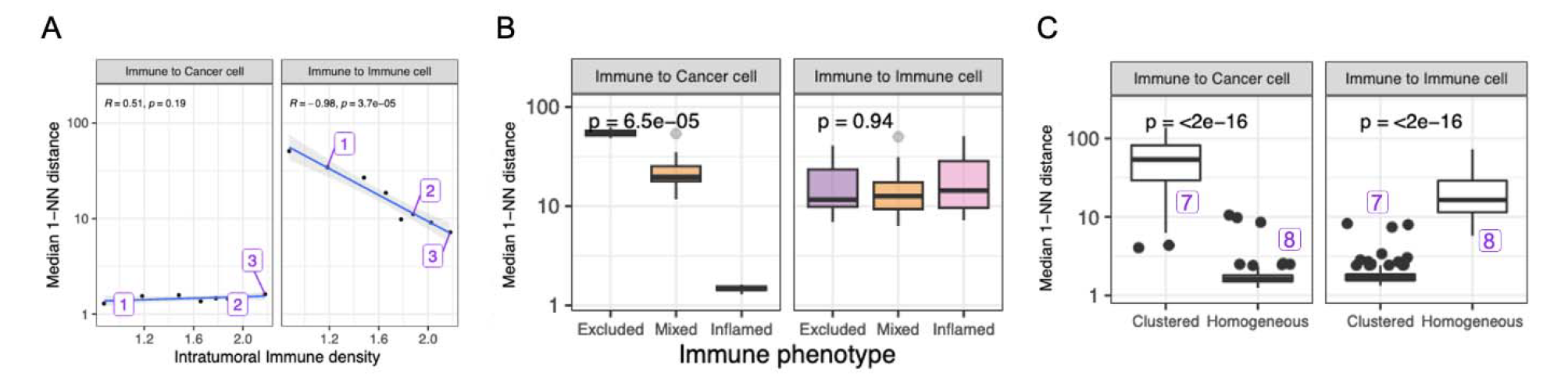
Associations between simulation study perturbations and spatial relationship parameters for a representative sample. (A) Median 1-NN distances vs. simulated intratumoral immune cell density for the SRs from Immune to Cancer cells and from Immune to Immune cell. Two-sided Pearson’s moment correlation test was used to test for the association. The Pearson’s coefficient and correlation p-value are represented in the plot. 1, 2, and 3 annotations match with Low, Medium and High densities from Supplementary Figure 9A. (B) Median 1-NN distances vs simulated immune phenotypes for the SRs from Immune to Cancer cells and from Immune to Immune cell. Differences between groups were tested by a Kruskan-Wallis test. (C) Median 1-NN distances vs simulated immune cell clustering for the SRs from Immune to Cancer cells and from Immune to Immune cell. A two-sided t-test tested differences between groups. All statistical tests were two-sided. Unless otherwise stated, no adjustments for multiple hypothesis testing were made. Abbreviations: 1-NN: first-nearest neighbor; SR: spatial relationship.

**Supplementary Notes Figure 3.**
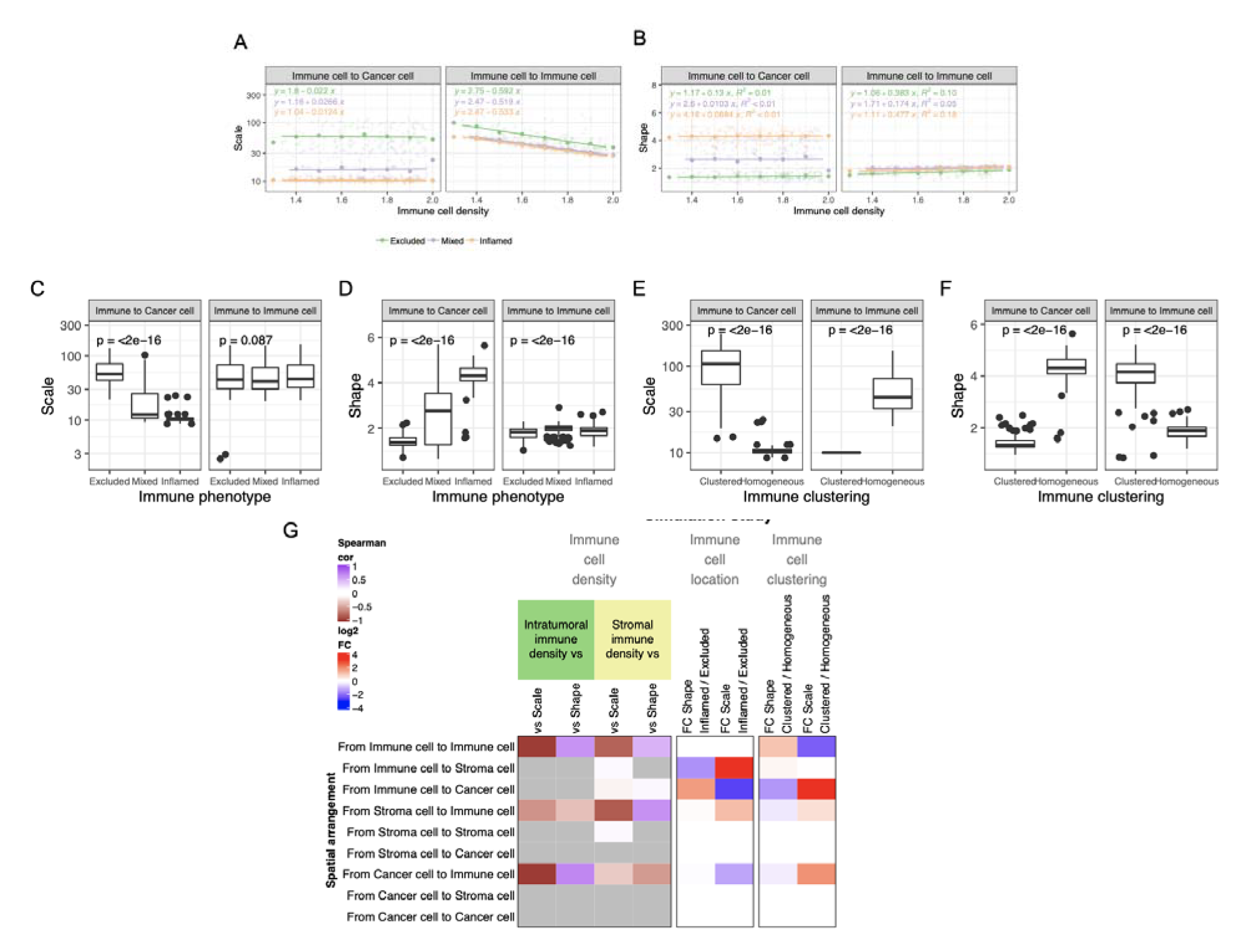
Associations between simulation study perturbations and spatial relationship parameters for the whole simulated cohort. (A) Scatter plot of the scale parameter for two representative SRs for simulations of immune cell at different density values (x-axis) and at distinct tissue compartments, including in the Tumor (Inflamed, orange), Stroma (Excluded, green) and in the Tumor and Stroma (Mixed, purple). A linear model was fitted using the data simulated at each tissue compartment (e.g. Inflamed), and the slope of the fit and significance is highlighted in the figure. Dots represent all simulations’ average scale parameters at a discretized density. (B) Scatter plot of the shape parameter for two representatives SRs for simulations of immune cell at different density values (x-axis) and at distinct tissue compartments, including in the Tumor (Inflamed, orange), Stroma (Excluded, green) and in the Tumor and Stroma (Mixed, purple). A linear model was fitted using the data simulated at each tissue compartment (e.g. Inflamed), and the slope of the fit and significance is highlighted in the figure. Dots represent all simulations’ average shape parameters at a discretized density. (C) Scale parameter distribution for simulations of immune cells present at different tissue compartments (Inflamed: in Tumor; Excluded: in Stroma; Mixed: in Tumor and Stroma). Each boxplot (e.g. Tumor) contains data for different samples and simulations at distinct densities. Statistical association was assessed by an ANOVA test. (D) Shape parameter distribution for simulations of immune cells present at different tissue compartments (Inflamed: in Tumor; Excluded: in Stroma; Mixed: in Tumor and Stroma). Each boxplot (e.g. Tumor) contains data for different samples and simulations at distinct densities. An ANOVA test assessed statistical association. (E) Scale parameter distribution for simulations of immune cells following a homogeneous spatial distribution, or a clustered spatial distribution. Each boxplot (e.g. clustered) contains data for different samples and simulations at distinct densities. A t-test assessed statistical association. (F) Shape parameter distribution for simulations of immune cells following a homogeneous spatial distribution, or a clustered spatial distribution. A t-test assessed statistical association. (G) First 4 columns: Spearman correlation between the spatial parameters (shape or scale) and intratumoral (green columns) or stromal immune cell density (yellow) for each SR. Fifth and sixth columns: fold change on differences on the shape or the scale parameters between Inflamed and Excluded simulations (non-significant fold changes are coloured in white). Statistical significance was assessed by a t-test. Seventh and eighth columns: fold change on differences on the shape and scale parameters between immune cell clustered or homogeneous simulations by a t-test. Non-significant correlations or associations are labeled in gray and white. Unless otherwise all statistical tests were two-sided, and no adjustments were made for multiple hypothesis testing correction. Abbreviations: *UC*: urothelial cancer; *SR:* Spatial relationship.

**Supplementary Notes Figure 4.**
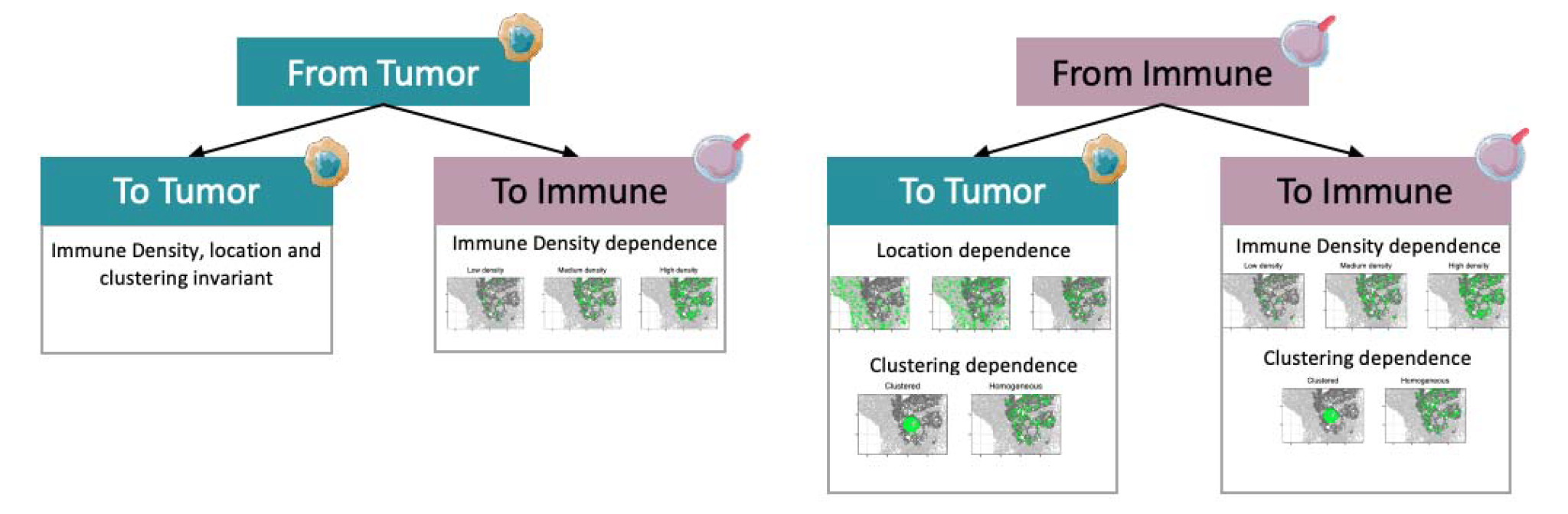
Interpretations on the associations between simulation study perturbations and spatial relationship parameters for the whole simulated cohort. Summary of associations between simulated perturbations and effect on SR parameters by reference and target cell type by aggregating data for the whole cohort (Supplementary Figure 21H). Here, the interpretation of when *Cancer cells* are replaced by *Negative cells* are analogous (abundant cell types). Immune cells are referenced in text text as rare cell types. Abbreviations: *SR*: spatial relationship.

## Extended Methods

### Spatial analysis: a simulation study to quantify spatial relationship parameters sources of variation

Using the segmentation between the tumor and stroma compartments and the tissue architecture, we simulated different states of immune cell infiltration or cell location to quantify sources of variation in the spatial patterns. We used our original samples’ point patterns and the delineated tumor and stroma compartments in all the simulations. We altered the abundance and location of three cell types: immune, cancer, and stroma.

First, we simulated immune cell density at different values. Here, all the cells belonging to the tumor compartment were labeled as cancer cells, and the cells belonging to the stroma compartment were labeled as stroma cells. Then, we randomly re-labeled cells a fraction of cancer cells (at a 1%, 2%, 5%, 7%, and 10% fraction) as immune cells, in which the SR of the immune cells resembled a homogeneous spatial distribution. Analogously, the same simulation study was carried out by randomly re-labeling stroma cells or altogether re-labeling cancer and stroma cells. These perturbations led to three different simulation studies (immune cells only in tumor, only in stroma, or in tumor and stroma), in which the SR parameters from/to cancer, immune and stromal cells (3×3=9 SRs) were estimated. For each simulation, the immune cell density (either intratumoral or stromal) was associated with the distinct SR Weibull parameters (shape and scale) obtained for the 9 SRs being studied.

In a second simulation study, we compared the SR parameters between the previous three different simulations that matched distinct immune phenotypes, being *Inflamed* (simulated immune cells in Tumor), *Excluded* (simulated immune cells in Stroma), and *Mixed* (simulated immune cells altogether in Tumor and Stroma). This study aggregated and compared simulations between groups, e.g., *Inflamed* vs *Excluded (*i.e., *Inflamed* combines simulations with an inflamed phenotype at multiple densities), as we aimed to quantify the global effect of the perturbation.

Lastly, we simulated differences in the local immune cell arrangement by perturbing the clustering of immune cells. In *Clustered* simulations, we allocated the immune cells next to each other by setting an anchor point in which immune cells were present at different abundances (1%, 2%, 5%, and 10% fraction of cells being immune cells). Using this simulation (*Clustered*), we compared the SR parameters with the previous simulation, in which immune cells were placed along the tissue following a *Homogeneous* spatial distribution.

